# Identifying unmeasured heterogeneity in microbiome data via quantile thresholding (QuanT)

**DOI:** 10.1101/2024.08.16.608281

**Authors:** Jiuyao Lu, Glen A. Satten, Katie A. Meyer, Lenore J. Launer, Wodan Ling, Ni Zhao

## Abstract

**Background:** Microbiome data, like other high-throughput data, suffer from technical heterogeneity stemming from differential experimental designs and processing. In addition to measured artifacts such as batch effects, there is heterogeneity due to unknown or unmeasured factors, which lead to spurious conclusions if unaccounted for. With the advent of large-scale multi-center microbiome studies and the increasing availability of public datasets, this issue becomes more pronounced. Current approaches for addressing unmeasured heterogeneity in high-throughput data were developed for microarray and/or RNA sequencing data. They cannot accommodate the unique characteristics of microbiome data such as sparsity and over-dispersion.

**Results:** Here, we introduce Quantile Thresholding (QuanT), a novel non-parametric approach for identifying unmeasured heterogeneity tailored to microbiome data. QuanT applies quantile regression across multiple quantile levels to threshold the microbiome abundance data and uncovers latent heterogeneity using thresholded binary residual matrices. We validated QuanT using both synthetic and real microbiome datasets, demonstrating its superiority in capturing and mitigating heterogeneity and improving the accuracy of downstream analyses, such as prediction analysis, differential abundance tests, and community-level diversity evaluations.

**Conclusions:** We present QuanT, a novel tool for comprehensive identification of unmeasured heterogeneity in microbiome data. QuanT’s distinct non-parametric method markedly enhances downstream analyses, serving as a valuable tool for data integration and comprehensive analysis in microbiome research.

## Background

Next-generation sequencing technology has facilitated large-scale microbiome profiling to explore associations with various disease and health outcomes. Given the large scale, samples must be collected and profiled in different runs or even from various studies. Furthermore, there has been a significant increase in the availability of publicly accessible microbiome data^1-3^, and integrating these diverse datasets can be invaluable in advancing scientific discoveries^4, 5^. When integrating these diverse datasets, it is crucial for researchers to recognize potential variations within these datasets, which might not be immediately apparent to external researchers. Much of this unwanted variation is attributable to hidden technical or biomedical heterogeneity, such as instrumental drift, undocumented differences in study design, or even lab labeling errors. Failing to account for these hidden heterogeneities can obscure the true biological signals and lead to spurious findings. The situation can be even worse when including publicly available data^1-3^, because sources of variability such as batch labels may not be included in the distribution data.

In genomic studies, numerous statistical methods have been developed to correct for batch effects when batch indicators are available, with ComBat being among the most widely adopted. In the microbiome field, several methods have also been proposed to address batch effects when indicators are known, including approaches such as MMUPHin^6^ and DEBIAS-M^7^ and ConQuR^8^, among others. These approaches rely on the availability of batch indicators. In contrast, no methods are designed to address *unmeasured/unknown* heterogeneity in microbiome studies, although techniques developed for genomic data, such as surrogate variable analysis (SVA)^9^ and remove unwanted variation (RUV)^10^, may be applicable. SVA is a two-step procedure for identifying surrogate variables (detection of hidden factors and then construction of surrogate variables), a general framework that was adapted by subsequent methods, including ours here. RUV employs factor analysis on control genes (via spike-in experiment or housekeeping gene estimation) to identify hidden heterogeneity. However, SVA and RUV are not ideally suited for microbiome data. First, they do not address the large number of zeros and the over-dispersion in microbiome data. Second, they only detect the underlying structure in the (possibly log-transformed) mean expression while ignoring other possible changes in distribution. Third, RUV requires users to specify the number of hidden directions: incorrect specification of these parameters may result in false positive or negative discoveries. Here, we present the Quantile Thresholding (QuanT) approach, the first tool for comprehensive identification of unmeasured heterogeneity in microbiome data. QuanT applies quantile regression^11^ at multiple quantile levels to taxa relative abundances, and then forms dichotomized residuals. QuanT distills information from these dichotomized residuals to output a set of unit vectors that represent the hidden heterogeneity. As a non-parametric approach, QuanT is robust to the complex distributional attributes of microbiome data, handles sparseness in a natural way, and can detect heterogeneity beyond simple mean shifts. The quantile surrogate variables (QSVs) output by QuanT can be easily incorporated into downstream analysis. QSVs can be treated as known variables and embedded into any existing analytic workflows (e.g., regression models) that allow for covariate adjustment. Alternatively, QuanT can be combined with the recently developed microbiome batch effect removal method, ConQuR^8^, to produce batch-free microbial profiles for downstream analysis.

## Methods

### Overview of QuanT

We first give an overview of the method before delving into a detailed explanation of each step. We propose a two-stage algorithm, depicted in Fig. 1, to identify unmeasured heterogeneity in microbiome relative abundance. Similar to existing methods such as SVA, our algorithm, QuanT, first employs matrix decomposition to detect hidden directions pertaining to unmeasured factors in the data, followed by a second stage to refine these detected directions. In the first stage, we fit quantile regression models at multiple quantile levels to the relative abundance of each taxon with the primary variable of interest and important covariates, and then generate dichotomized, centered quantile residuals. These columns of residuals are concatenated into a single matrix of discretized residuals *RZ* We then conduct a singular value decomposition (SVD) of *R* to uncover the systematic structure of the residuals and select singular vectors based on their directional information and variability. These selected vectors, termed stage-1 singular vectors, capture systematic variation independent of the known variables.

**Fig. 1.**
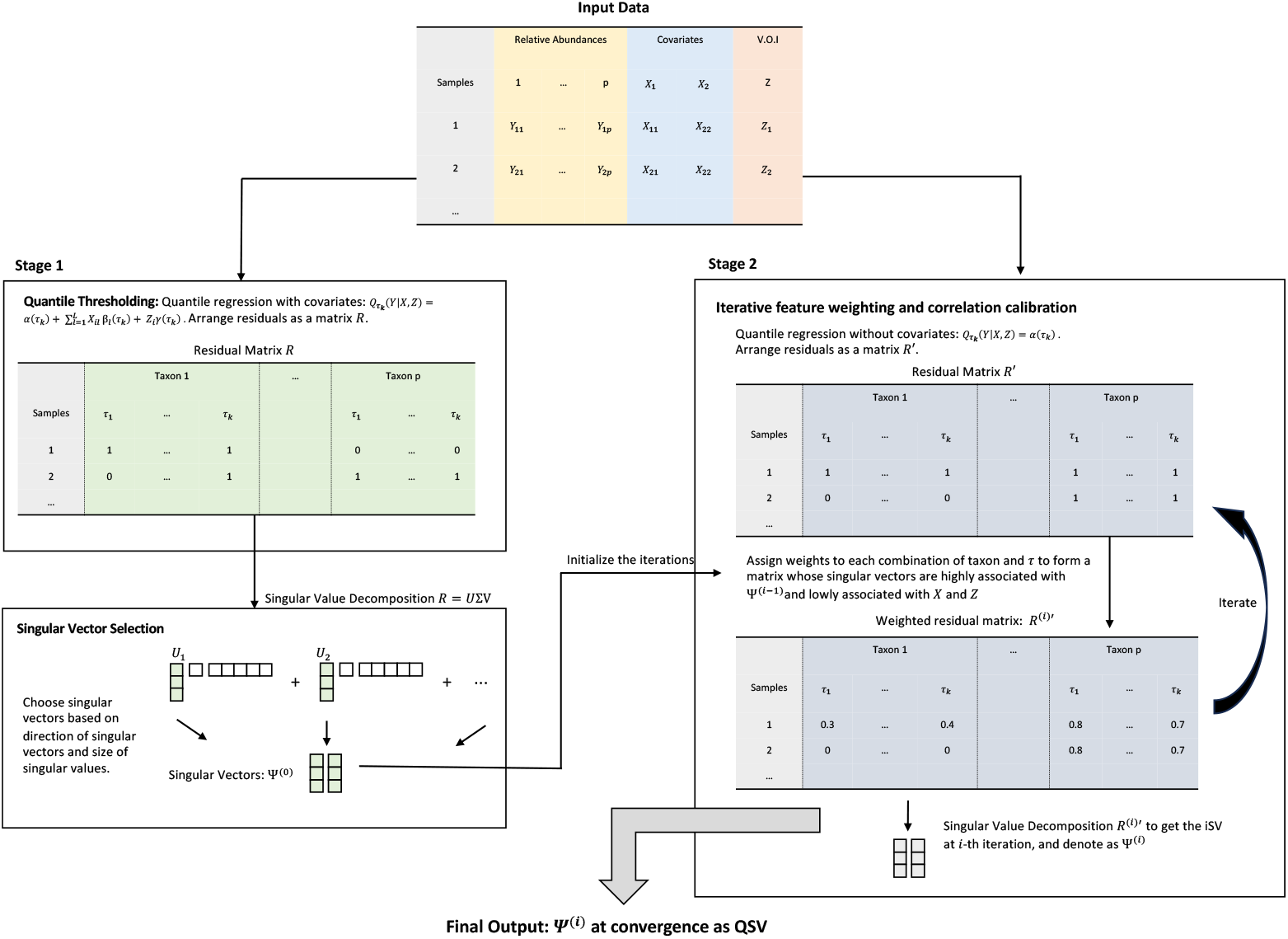
Flowchart of QuanT. V.O.I: variable of interest; QSV: quantile surrogate variable; τ_*i*_ … τ_*j*_: quantile levels. Stage 1 conducts quantile regression for each taxon at multiple quantile levels using V.O.I (*Z*) and covariates (*X*). Residuals are thresholded to create a binary matrix *R*, which is analyzed through singular value decomposition to identify key directions. Stage 2 implements iterative feature weighting and correlation calibration. Stage 2 starts by fitting quantile regression and thresholding without covariates at multiple quantile levels, followed by the weighting of taxon and quantile levels so that the singular vectors of the weighted residual matrix *R*^(*u*)′^ are highly associated with the prior stage singular vectors but minimally associated with *X* and *Z*. The singular vectors at convergence of Stage 2 are QSVs, the output of QuanT.

The stage-1 singular vectors are unrelated to the known variables as these known variables were included as covariates in the quantile regression models. As a result, adjusting for stage-1 singular vectors in downstream analysis would not correct for the confounding effects of the unmeasured heterogeneity (although efficiency may be improved). Therefore, in the second stage, we perform a “correlation calibration” step, in which we weight the “features” (i.e., combinations of quantile level and taxon). Specifically, we upweight features that are strongly marginally associated with the stage-1 singular vectors but weakly associated with the covariates, and downweight those with the opposite characteristics. Our second stage is an iterative procedure such that in each iteration, we fit quantile regression models and decompose a matrix of discretized residuals, similar to the first stage, but using the weighted features for model fitting. Each iteration will yield new “iterative” singular vectors (iSVs) by decomposing the residual matrix. The iSVs at convergence, i.e., when the resulting iSVs are close enough to the iSVs in the previous iteration, are called Quantile Surrogate Variables (QSVs), which are the output of our algorithm. This kind of correlation calibration step has been shown to be necessary in similar tasks when analyzing gene expression data^9, 12-15^.

Comparing to the existing approaches such as SVA and RUV, QuanT offers several novel modifications/improvements to make it suitable for microbiome data, which is typically sparser and over-dispersed. First, by using quantile regressions instead of least squares or generalized linear models, QuanT can gather information from the whole distribution of the taxa rather than just the mean, which is critical for microbiome data. Secondly, SVA and RUV search for significant directions by identifying singular vectors that represent a disproportionately high amount of variation. QuanT, on the other hand, relies on both the direction of a singular vector and the variance it explains, and thus allows us to detect significant directions that may harbor important underlying structure but explain only a modest proportion of variation.

We now describe our notation and proceed to a more in-depth explanation of the method’s components.

### Notation

Assume the relative abundance data form a *n* × *p* matrix *Y*, in which *Y*_*ij*_ is the relative abundance of the *j* -th taxon in the *i* -th sample. We denote by *Y*_.*j*_ the *j*-th column of *Y* comprising data on the *j*-th taxon, and treat it as the outcome in the proposed regression framework. Suppose that in addition each sample *i* has a primary variable of interest (denoted by *Z*_*i*_) and a set of *L* covariates (denoted by *X*_*i*_) to adjust for. We use quantile regression at each of *K* quantile levels in our regression framework; these levels are represented by the vector,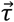. By default, we set the quantile levels as an equally spaced sequence from 0.1 to 0.9 with a step size of 0.005 corresponding to K=161.

Note that some taxon-quantile combinations may not convey useful information. For example, if 70% of the observations for a specific taxon *j* are zeros, any quantile levels 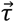 less than 70% will be irrelevant and are thus disregarded for this taxon. Thus, the effective length of 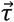 varies by taxon. This approach makes 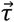 data adaptive, utilizing more quantile levels for taxa with more informative distributions.

### Stage 1: Obtaining singular vector directions that capture the underlying heterogeneity structure

#### Quantile thresholding

For each feasible pair of taxon *j* and quantile level τ_*k*_, we fit the quantile regression Model 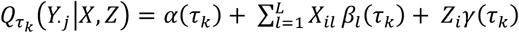 to the data *Y*_.*j*_ in which 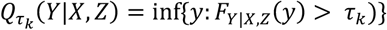 is the τ_*k*_ -th quantile function. Parameters 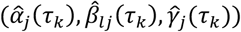 are obtained by solving the minimization problem

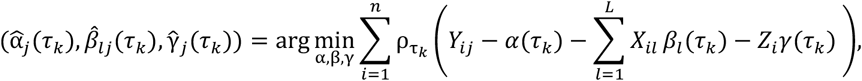

where 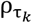 is the check function^16^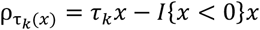.

After fitting the quantile regression model, we threshold the microbial abundances against the estimated quantiles and center them as 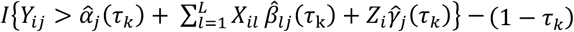. For each pair of taxon *j* and quantile level τ_*k*_, this yields a *n*-dimensional vector of discretized (binary) centered residuals. We gather the results from different pairs into a matrix *R* with *n* rows (samples) and no more than *pK* columns (processed features) as we omit the uninformative pairs of taxon and quantile level. *R* captures the heterogeneity structure such as the residual matrix constructed by SVA but is more robust and distribution-free and handles zero cells in a natural way.

#### Singular vector selection

We conduct SVD of the binarized and centered residual matrix *R* to obtain

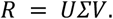

We aim at selecting a subset of the singular vectors, i.e., columns of *U*, to represent the important structure in the data. We use an integrated approach to select the singular vectors. Specifically, we select singular vectors 1) whose distributions are statistically different from the marginal distribution of the singular vectors if there is no underlying structure in the binary centered residual matrix (a.k.a., complete null, details are discussed in the next paragraph), and which explain a large proportion of variability of the binary centered residual matrix. We use the SelectiveSeqStep procedure in the knockoff paper^17^ to achieve this goal based on the two criteria simultaneously. This dual goal selection is a novel feature of QuanT. The detailed rationale and procedure are as follows.

Under the complete null that there is no underlying structure in the binary centered residual matrix, each singular vector direction should point to a uniformly distributed point on the *n-*dimensional sphere; this implies that the components of each singular vector should follow a Gaussian distribution^18^. Thus, we can test whether a singular vector is an important direction by testing the normality of each singular vector direction using the Lilliefors test^19^ or the Anderson-Darling test^20^ or the Shapiro-Francia test^21^ and obtain its corresponding p-value. We also order the singular vectors based on the size of their corresponding singular values. We aim to select from the top *m* singular vectors which have p-values smaller than a cutoff *c* (e.g., 0.05). The choice of *m* is determined by controlling the false discovery rate (denoted by *q*) of the selected singular vectors. Under the null hypothesis, p-value is uniformly distributed; thus the ratio between the number of singular vectors with p-values greater than *c* and those with p-values less than *c* should be 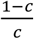. Roughly speaking, the singular vectors with p-values > *c* can be considered generally null, and therefore, we can estimate the number of falsely discovered singular vectors based on them, which is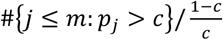, where # represents the size of a set. This is similar to the idea of q-value^22-24^, a commonly used approach for multiple testing problems. The proportion of falsely discovered singular vectors, if we select the top *m* singular vectors, are estimated by 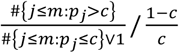. We pick the *m* such that this proportion is controlled below a false discovery rate threshold *q* if the top *m* singular vectors are selected and would exceed *q* if the top *m* + 1 singular vectors were selected:

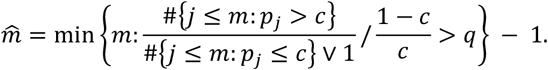

After determining 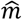, we select the singular vectors from the top 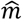 singular vectors with p-values smaller than *c*. Note that our approach allows for a non-monotonic selection of singular vectors, yet primarily focus on those that account for a substantial portion of variability in the data.

We define Φ to be the matrix with the *s*-th column, denoted as Φ_*s*_, equal to the *s-*th stage-1 singular vector.

### Stage 2: Iterative feature weighting and correlation calibration

Similar to SVA^9, 12-15^, we conduct a second stage to refine the stage-1 singular vectors that we call correlation calibration. The reason is that, by regressing out *X* and *Z* to construct the binarized centered residuals in the quantile regression stage, the selected singular vectors are, loosely speaking, uncorrelated with the primary variable *Z* and covariates *X* (if we had used linear regression, this statement would be exact). Thus, they are not able to capture unmeasured confounding effects, as confounders must be correlated with *Z* as well. Nonetheless, it is the unmeasured confounders that pose the greatest threat to the integrity of a study.

Because the stage-1 singular vectors represent the important features of the full set of residuals *R*, we seek to reweight the columns of *R* that will allow us to identify directions that not only reflect the underlying data structure but also maintain potential correlations with *X* and *Z*, thereby enhancing the robustness of downstream analysis against hidden confounding factors.

To accomplish this, we first conduct a similar thresholding procedure as in stage 1, but on a reduced quantile regression model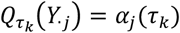, i.e., without regressing *X* and *Z* out, and construct the binarized and centered residuals (denoted by *R*^′^). We then weight columns (combinations of taxon and quantile levels) of *R*^′^ to prioritize those that represent underlying data structure while being minimally associated with *X* and *Z*. Specifically, we use an iterative procedure that balances the two goals—capturing the underlying heterogeneity and maintaining appropriate correlation with *X* and *Z*.

In the first iteration, for each feasible pair of taxon *j* and quantile level τ_*k*_, we fit two pairs of quantile regression models to the columns of *R*^′^. The first pair compares the reduced quantile regression model 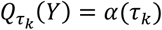to the model that includes the stage-1 singular vectors 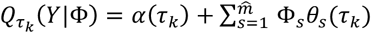. The second pair compares the model with the stage-1 singular vectors 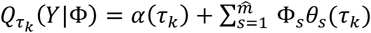 to the full model 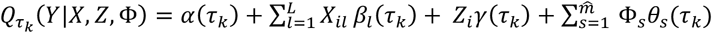. Note that the two models in the first pair can assess whether the taxon-quantile combination can capture information in the stage-1 singular vectors, and the second pair of models allows us to assess whether the taxon-quantile combination convey information about the known covariates, conditional on the stage-1 singular vectors. Thus, these two pairs of models facilitate the identification of directions that relate to the stage-1 singular vectors without being overly influenced by *X* and *Z*.

For each pair of hypotheses at each taxon *j* and quantile level τ_*k*_, we calculate a rank-based pseudo-F statistic^25^ to generate a p-value, comparing whether the larger model fits the data better than the smaller model. With this, we obtain two sets of p-values, with 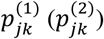 corresponding to the p-value comparing the first (second) pair of models for taxon *j* and quantile level τ_*k*_. Intuitively, a small 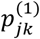 implies that the stage-1 singular vectors corresponding to taxon *j* and quantile level τ_*k*_ convey important information on the data structure; yet a large 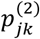 suggests that taxon *j* and quantile level τ_*k*_ is less impacted by the known covariates given the corresponding stage-1 singular vectors. Thus, we will weight each column of *R*^′^ so that columns with small 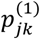 values and large 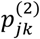 values receive a higher weight.

To be specific, we apply the local FDR method^26^ to the two sets of p-values separately to obtain two sets of local FDR estimates. This transforms each set of p-values to a set of local FDR estimates. Each taxon-quantile pair corresponds to two p-values before the transformation, and hence corresponds to two local FDR estimates after the transformation.

These two local FDR estimates answer the following questions: given the observed data for this specific taxon-quantile pair, what is the posterior probability that it is impacted by the unmeasured heterogeneity and what is the posterior probability that it is not impacted by the known covariates. We then use the product of the two local FDR estimates as the weight for the taxon-quantile pair. We multiply each column of *R*′ by its weight and obtain a reweighted matrix *R*^(*u*)^′. We then conduct SVD to obtain a new set of singular vectors, which we denote as Ψ^(1)^, in which the superscripts indicate the iteration cycle, and we are now at iteration 1.

Note that *R*^′^ is obtained without regressing out *X* and *Z*, and thus, Ψ^(1)^ can be correlated with *X* and *Z*. Note also that the number of the new singular vectors Ψ^(1)^ can be larger than the stage-1 singular vectors Φ. We seek to only maintain the directions that are most correlated with Φ. To do this, for each column, Φ_s_, we search for the columns of Ψ^(1)^ that have the highest absolute correlation with Φ_s_. In other words, for each column Φ_*s*_, we choose the *d*_*s*_-th column of Ψ^(1)^ in which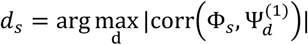. The final selected singular vectors are called the iSVs at the first iteration.

For later iterations, we replace the stage-1 singular vectors in the two pairs of regression models with the iSV of the previous iteration and follow the same procedure. The iteration stops when iSVs from two adjacent iterations are the close enough, or when we reach a sufficient number (*T*_*max*_, with default = 10) of iterations. In the second case, we calculate the final singular vector as a weighted average of the last *T*_*max*_/2 iterations. The final singular vectors are the QSV output of our algorithm. Please refer to the Algorithm 1 for individual steps.

#### Algorithm 1: QuanT algorithm

**Input: Y, X, Z**

**Output: Ψ (quantile surrogate variables)**

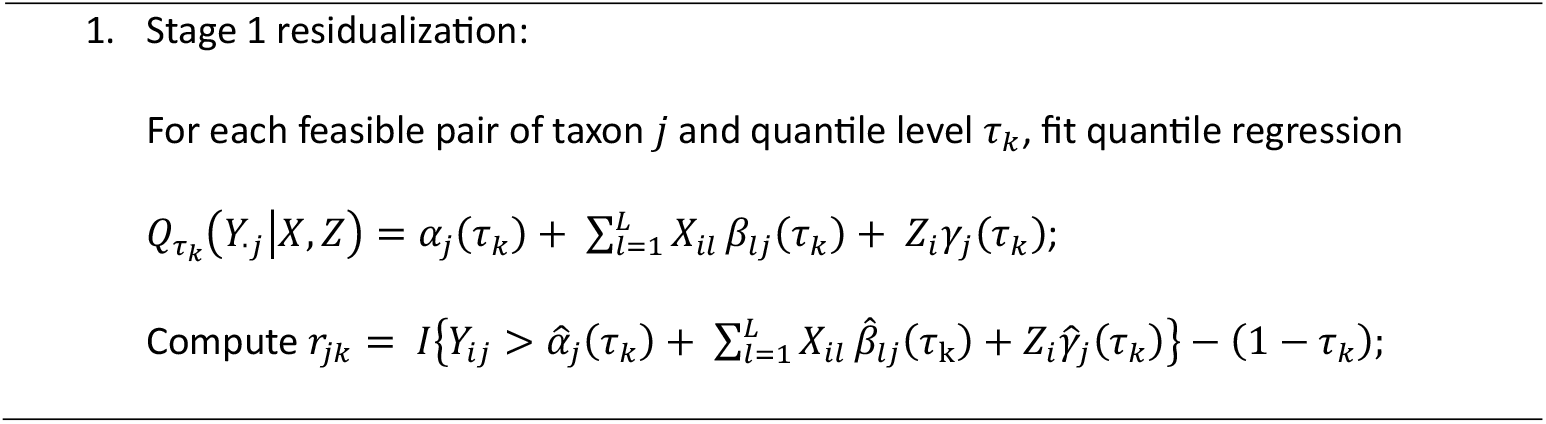
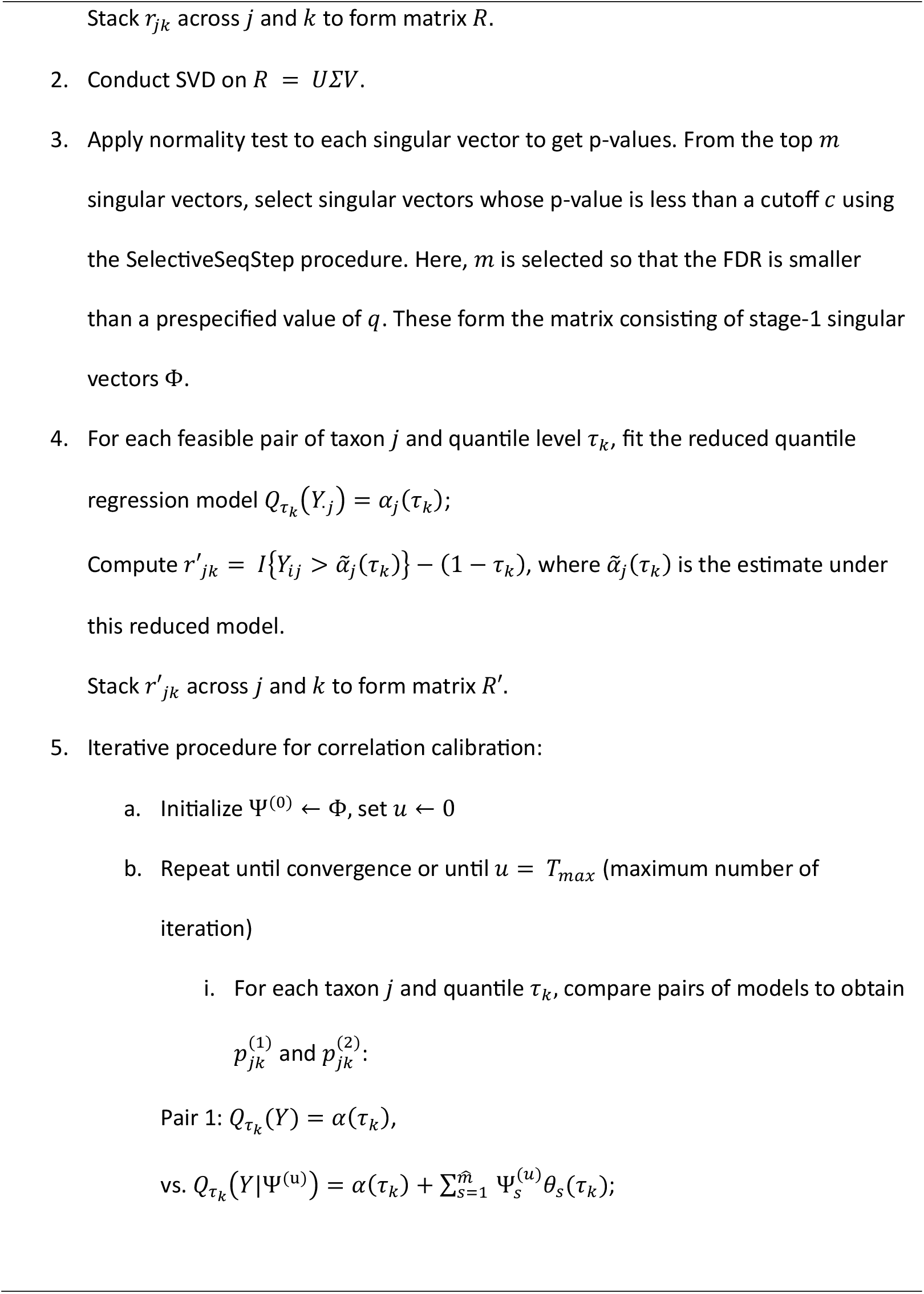
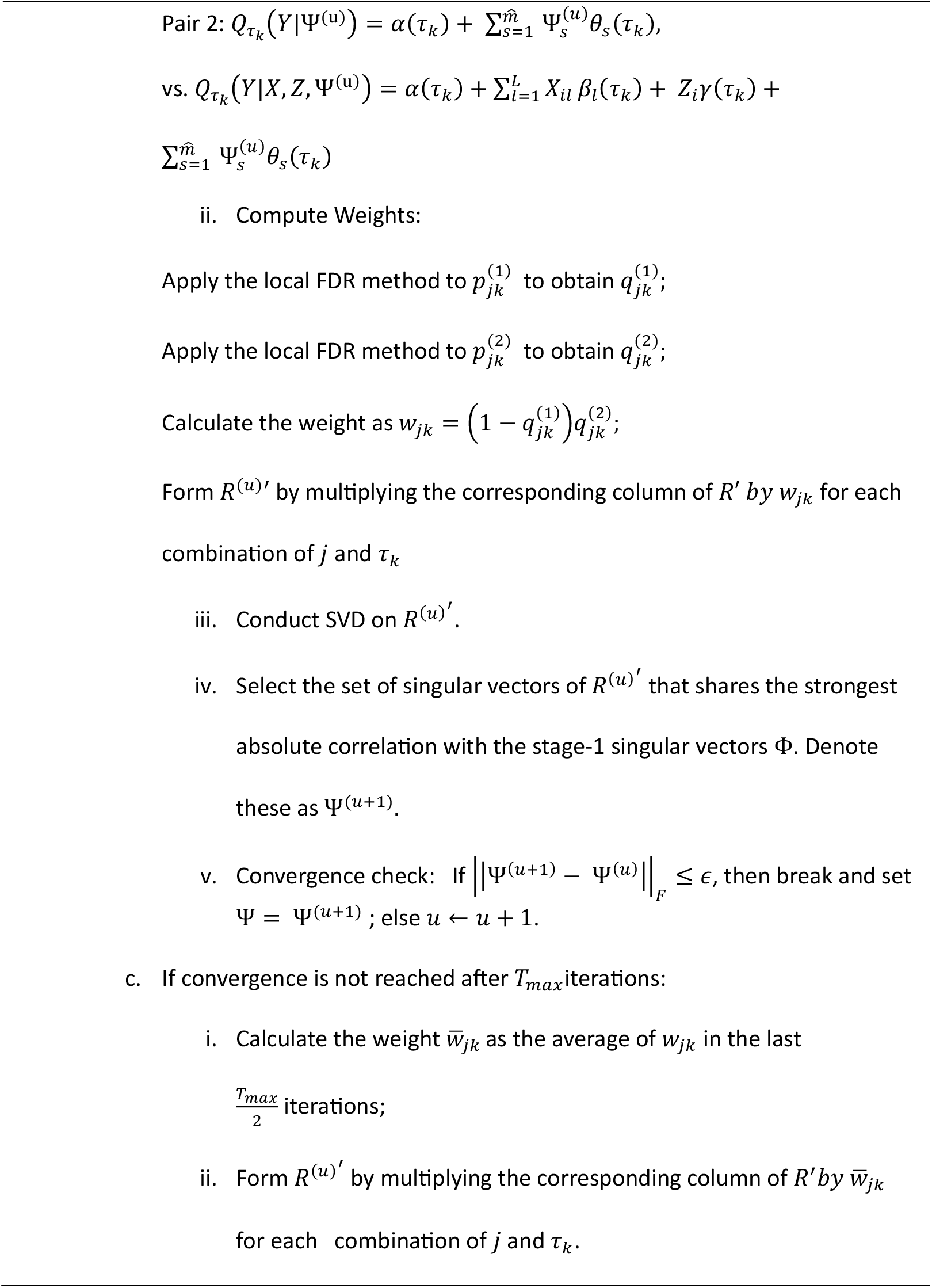
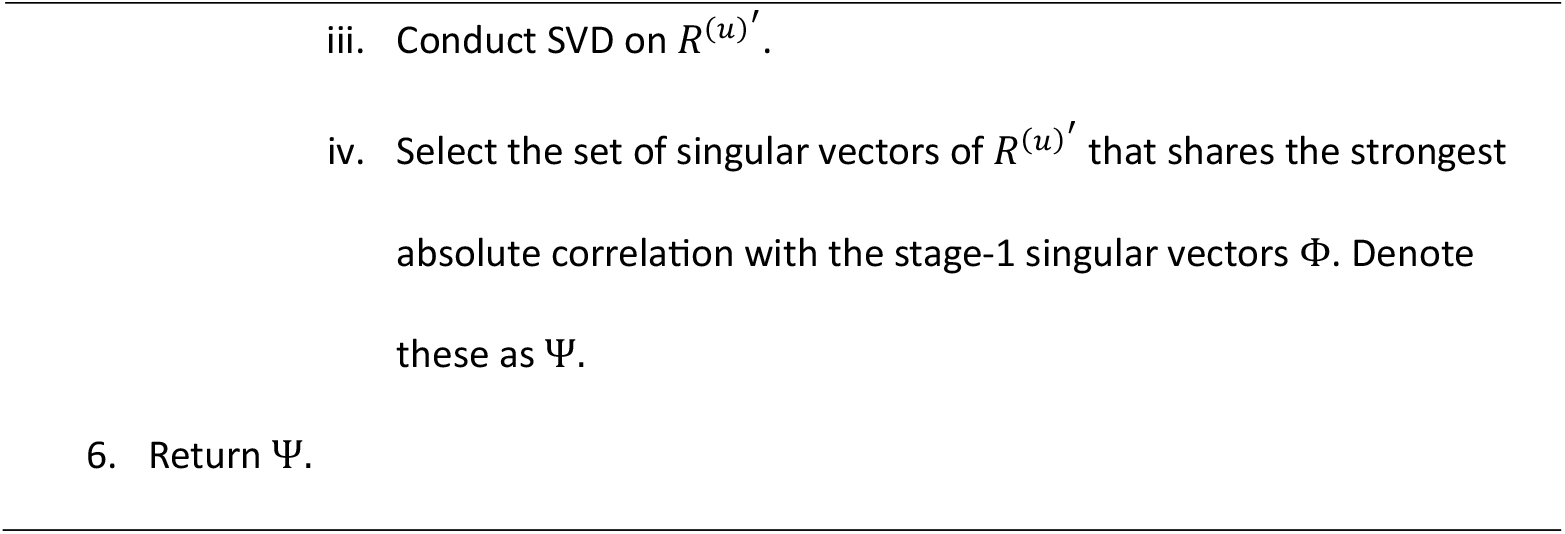

### Singular vectors selection when no stage-1 singular vectors are found

In rare cases, we may encounter the situation where no singular vector is selected in the first stage. This usually happens when the hidden heterogeneity variable is normally distributed or when the sample size is not large enough. In this case, we provide an alternative approach that selects important singular vectors solely based on the proportion of variabilities explained. This approach is very similar to the principal component/eigengene selection methods in SVA, except that the least squares regression models are replaced by appropriate quantile regressions. The detailed algorithm can be found in the Algorithm 2. Note that this variant in our algorithm only applies to calculating the stage-1 singular vectors; we still need the same iterative correlation calibration procedure as in the standard QuanT algorithm (Algorithm 1).

#### Algorithm 2 alternative stage-1 algorithm when no singular vector is selected through the normality test

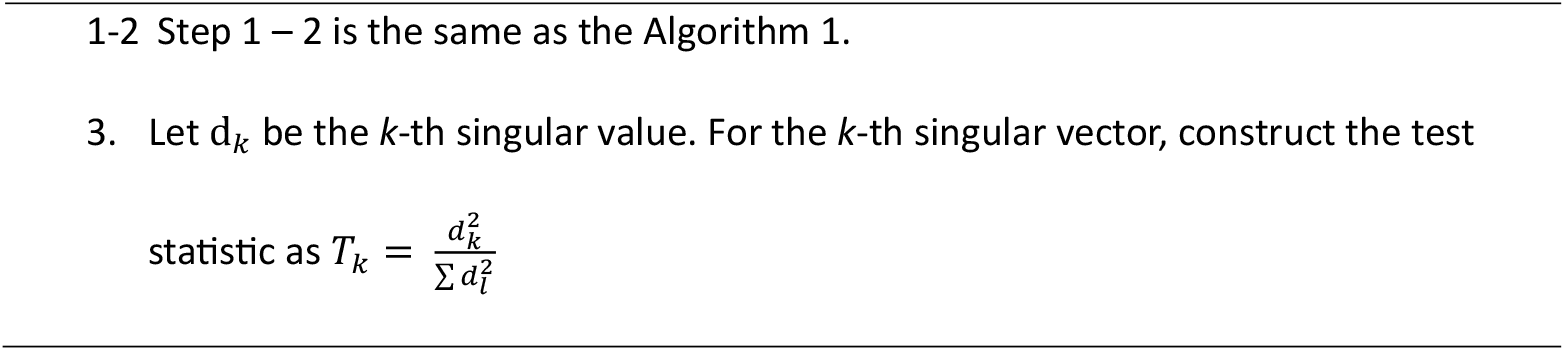
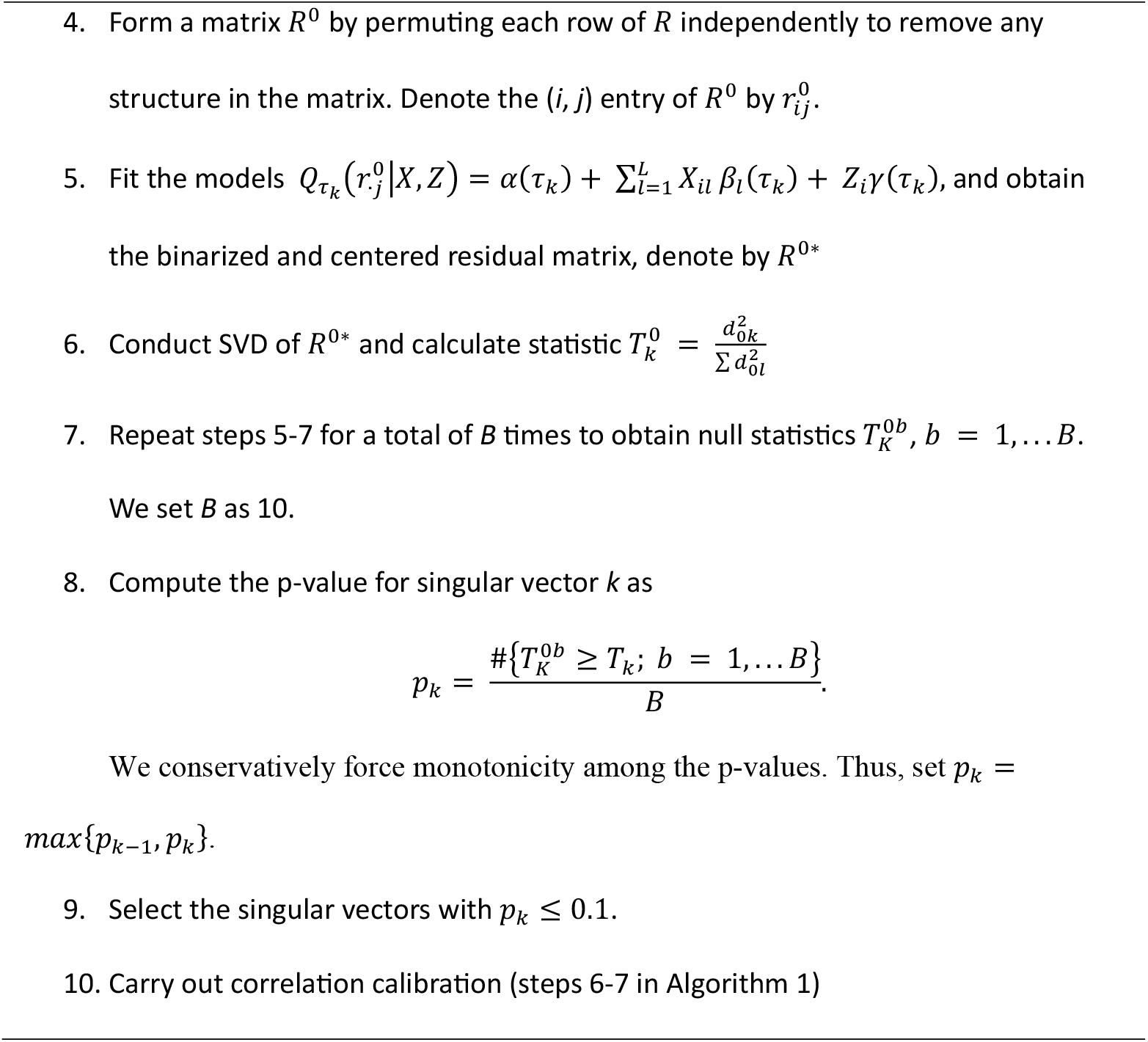

### Simulation settings

We conducted simulations to evaluate QuanT’s performance when the true signal is known. We simulated data that mimicked two datasets from the integrative Human Microbiome Project^27^: a gut microbiome dataset from the Inflammatory Bowel Disease Multi-omics Database (IBDMDB) project^28^, and a vaginal microbiome dataset from the Multi-Omic Microbiome Study: Pregnancy Initiative (MOMS-PI) project^29^. These two datasets represent microbiome data from two body sites that are of great research interest. They also have contrasting attributes in terms of sparsity and over-dispersion level, which makes them useful for evaluating our proposed method QuanT. We used two methods to simulate data that mimicked the template data: the parametric Dirichlet-Multinomial (DM) model and the non-parametric approach in MIDASim^30^. We additionally simulated a third scenario where microbiome absolute abundances were generated using a log-linear model, in order to evaluate QuanT’s performance in compositional settings.

For each sample *i*, we simulated a clinical variable of interest (denoted by *Z*_*i*_) and a hidden batch variable (denoted as *B*_*i*_) through a bivariate Bernoulli distribution, with both variables having a marginal mean of 0.5. A correlation of 0.3 was set between them. Note that because the “batch” variable is simulated to be correlated with both the microbiome data and *Z*, thus, it is a confounder. Here, we use the “batch” variable to represent all the hidden heterogeneities.

For the parametric simulations, we adopted the simulation setup from a previous study^31^ to simulate taxa abundances using the Dirichlet-Multinomial (DM) distribution. In our simulations, we designated the batch variable to impact most taxa (referred to as Batch Taxa), while the variable of interest, *Z*, influenced only a small subset (referred to as Clinical Taxa). Specifically, for the IBDMDB dataset, we randomly selected 250 of the 301 taxa to be Batch Taxa, among which 40 were also Clinical Taxa. Similarly, for the MOMS-PI dataset, we selected 200 of the 233 taxa as Batch Taxa, with 50 of these being Clinical Taxa. We first fitted the DM model to the template datasets (IBDMDB and MOMS-PI) and obtained the estimated mean vectors *π*_*0*_. Subsequently, we formed two additional vectors *π*_*B*_ and *π*_*Z*_ to incorporate batch and clinical effects. Both vectors were initially set to be the same as π_0_. We then randomly permuted the frequencies within *π*_*B*_ and *π*_*Z*_ corresponding to the Batch Taxa and Clinical Taxa. Finally, a sample-specific frequency was defined as *π*_*i*_ = (1 − *w*_*B*_*B*_*i*_ − *w*_*z*_*Z*_*i*_)*π*_*0*_ + *w*_*B*_*B*_*i*_*π*_*B*_ + *w*_*z*_*Z*_*i*_*π*_*z*_, where (*w*_*B*_, *w*_*z*_) determine the underlying batch and clinical effects. We considered three combinations for these parameters: (*w*_*B*_, *w*_*z*_) ∈ {(0,0*Z*5), (0*Z*5,0*Z*5), (0*Z*8,0*Z*2)}. When *w*_*B*_ = 0, there is no underlying hidden heterogeneity/batch effect. The count data for each sample were generated using the DM distribution with *π*_*i*_ and an overdispersion parameter of 0.02.

For the non-parametric simulations, we used the Microbiome Data Simulator (MIDASim)^30^, which is designed to generate microbiome data that capture the underlying distributional and correlation structure of a template dataset. It offers a nonparametric quantile matching approach, and a parametric approach based on log-linear models. Here, we utilized the non-parametric version because it captures the underlying data structure more accurately. Via MIDASim, for each template data set, we first generated vectors of relative abundances and zero-proportions. After choosing the Batch Taxa and the Clinical Taxa, we generated sample-specific relative abundance and zero-proportion vectors in the same way as in the DM simulation and also used the same (*w*_*B*_, *w*_*z*_) combinations. Count data for each sample were generated by inputting the sample-specific vectors into MIDASim and choosing to use its non-parametric approach.

In the third simulation scenario, we again simulated microbiome read counts using the Dirichlet−Multinomial model, but introduced batch and clinical effects differently. Instead of modifying the mean frequency vectors *π*_*B*_ and *π*_*Z*_ as in the first simulation, we first generated read counts from the fitted DM model using *π*_*0*_, and then directly adjusted the counts. Specifically, for samples with *B*_*i*_ = 1, read counts of Batch Taxa were multiplied by *w*_*B*_; for samples with *Z*_*i*_ = 1, read counts of Clinical Taxa were multiplied by *w*_*z*_. We considered three combinations for these parameters: (*w*_*B*_, *w*_*z*_) ∈ {(1,4), (4,4), (5,3)}.

## Results

To rigorously assess QuanT, we conducted extensive simulation studies based on real-world gut and virginal microbiome datasets from the HMP2 projects^27^. We also applied QuanT and competing methods to three curated microbiome datasets, each of which has distinct sources of technical heterogeneity: a gut microbiome study on cardiovascular diseases, which displays a moderate technical variation arising from sequencing in multiple runs; an integrated dataset compiled from various HIV gut microbiome studies; and a consolidated dataset encompassing multiple colon cancer studies. Through both visual and quantitative comparisons, we demonstrate that QuanT outperforms existing methods by robustly and comprehensively uncovering underlying microbiome data heterogeneity, which is beyond mere shifts in means. When incorporated into the analytical workflow, QuanT effectively reduces the influence of unwanted heterogeneity, enhancing false discovery rate control, maintaining sensitivity of discovery, as well as improving prediction accuracy.

### Application to the CARDIA study, a single large-scale epidemiological investigation

We assessed QuanT using multiple datasets. We first applied it to a study containing traditional batch variation: samples are collected under one protocol but handled in different batches. The Coronary Artery Risk Development in Young Adults (CARDIA) Study^32^, initiated in 1985-1986, recruited young adults aged between 18 and 30 with the goal of uncovering determinants of cardiovascular disease (CVD) throughout adult life. Subject enrollment was balanced according to race, gender, education, and age. Extensive data on risk factors related to CVD have been collected, including health behaviors, medical history (including medication use), and clinical risk factors, such as blood pressure. Stool samples were also collected for some visits. Here, we used the microbiome data generated from the stool samples collected during the Year 30 follow-up (2015-2016). The samples were processed through four DNA extraction/library preparation batches, and subsequently sequenced through seven runs using the Illumina MiSeq 2×300 platform. Specifically, each DNA extraction batch was allocated to two sequencing runs, except for the final batch, which was sequenced by itself during the seventh run. Consequently, the data were processed in seven distinct batches. Following sequencing, forward reads were processed through the DADA2^33^ pipeline for quality control and derivation of amplicon sequence variants (ASVs), and taxonomy was assigned using the Silva reference database^34^. The data were agglomerated to the genus level, and taxa that did not yield any reads in all samples were omitted from the dataset. After preprocessing, the data contains 633 samples and 145 taxa. Analyses were restricted to participants for whom all covariate data were available.

In this study, we considered high blood pressure (HBP) (non-HBP = 0, HBP = 1) as the variable of interest, and gender (Male = 0, Female = 1) and race (White = 0, Black = 1) as adjustment covariates. To assess QuanT and the competing methods, we treated the batch indicators as unknown and used these methods to identify the now-hidden heterogeneities/batches. The performance of these methods was assessed based on three criteria: 1) consistency and sensitivity in downstream differential abundance analysis for individual taxa; 2) enhancement in prediction accuracy for the variable of interest; and 3) the efficacy of batch effect removal while preserving the effect from the variable of interest at the entire community level. For SVA, we used the svaseq function, which added a pseudo count of 1 to all counts and then applied log transformation as described in Leek (2014)^15^, and used the iterative reweighted algorithm^12^. The default parameter setting was used. For RUV, we used the RUVg approach in Risso *et al*. (2014)^10^ with an estimated group of null taxa as controls, rather than RUVr and RUVs as these approaches can only identify hidden directions that are uncorrelated with the variables of interest. For hyperparameters, we selected control taxa using the Benjamini−Hochberg procedure on edgeR p-values at a 0.05 threshold, and set the number of latent factors following the guidance in Risso *et al*. (2014)^10^.

Fig. 2a and 2b compare the consistency of differential abundance analysis results across various models. Here, differential abundance analysis was conducted using linear regression on relative abundances of each taxon, controlling for age, gender, and identified hidden variables. For benchmarking purposes, analyses were conducted adjusting for true batch/group indicators (TG), which was regarded as the gold standard, and with no correction (NC), which served as the baseline. Fig. 2a shows the number of differentially abundant taxa identified from the CARDIA dataset by Benjamini-Hochberg adjusted p-value < 0.05. TG leads to 4 discoveries. Considering these as true signals, NC identifies the 4 true discoveries and has 1 additional false discovery; QuanT identifies 3 true discoveries and has 1 additional false discovery. SVA and RUV lead to no discoveries. Therefore, QuanT demonstrates superior performance compared to SVA and RUV when applied to the CARDIA dataset, identifying more potential true discoveries while maintaining a reasonable false discovery proportion. Fig. 2b shows the top K detected differentially abundant taxa. Treating TG as the gold standard and calculating the proportion of overlapping discoveries, QuanT has comparable performance with NC and SVA, and is substantially better than RUV.

**Fig. 2.**
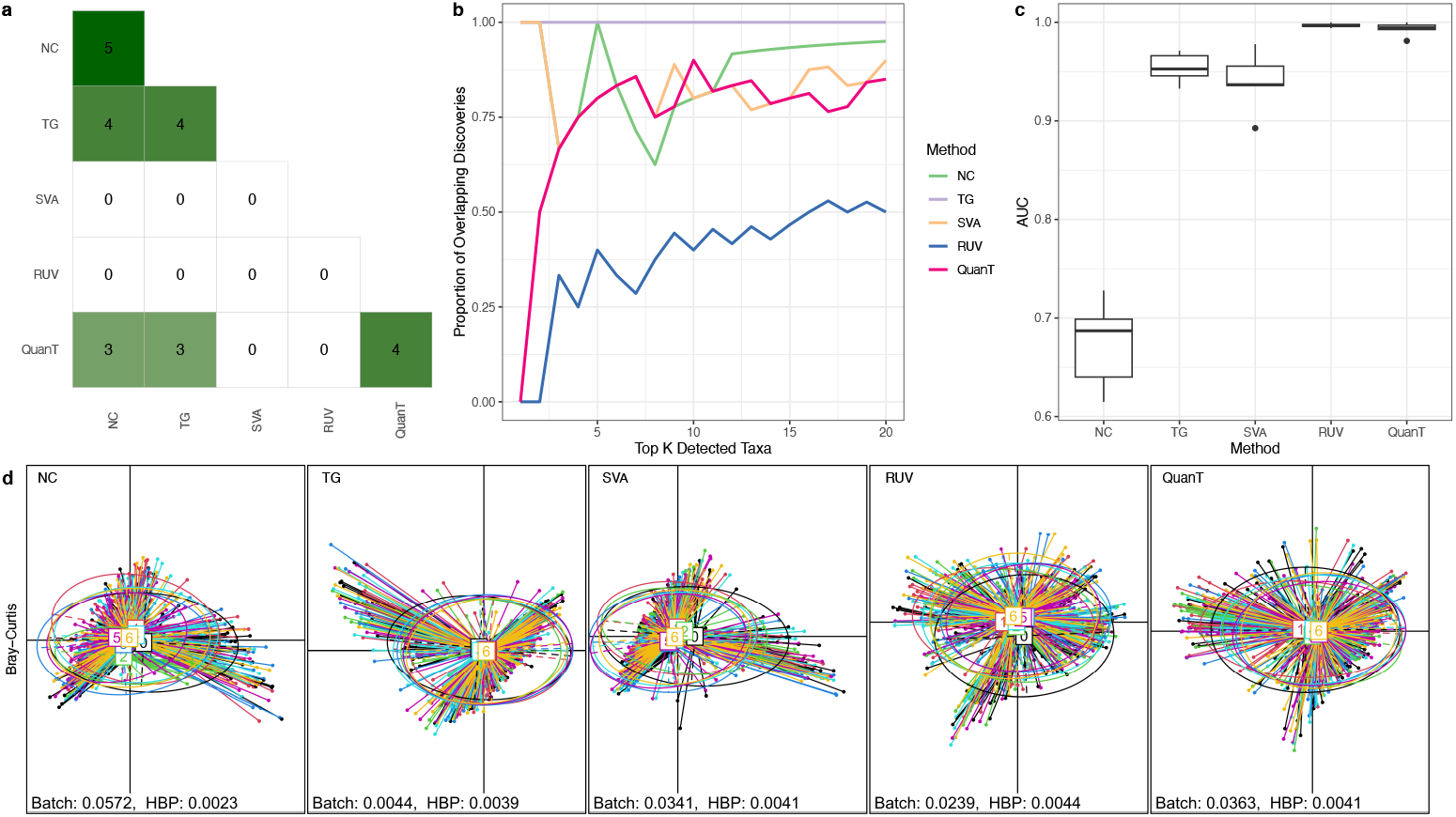
Evaluation on the CARDIA data. NC: no batch correction; TG: correction using true group indicators. **a** Heatmap representing the number of differentially abundant taxa identified in the CARDIA dataset by Benjamini-Hochberg adjusted p-value < 0.05; diagonal elements are numbers detected, and off-diagonal elements are numbers detected in common. **b** Proportions of overlapping taxa in the top K detected differentially abundant taxa between each method and TG in the CARDIA dataset. **c** Average cross-validated area under the receiver operating characteristic curves (ROC-AUC) of predicting HBP from the ConQuR-corrected microbiome data via random forest. Larger values indicate stronger predictive capabilities. **d** PCoA plots of the ConQuR-corrected data based on Bray-Curtis distance, with PERMANOVA R^2^ of batch ID and HBP.

Fig. 2c shows the prediction accuracy using the original and the ConQuR corrected microbiome data, where the corrected data were adjusted for TG, or surrogate variables estimated by SVA, RUV or QuanT. Random forest models were fit with five-fold cross-validation to predict HBP status, and the area under the receiver operating characteristic curve (ROC-AUC) was used to evaluate the accuracy. We note that this is not a genuine predictive analysis, but rather a metric to quantify how well the corrected microbiome data can explain the variable of interest. As shown in Fig. 2c, RUV and QuanT corrected data predicts HBP more accurately than using TG or SVA, and prediction using the unadjusted data (NC) leads to the worst accuracy. Fig. 2d shows the principal coordinate analysis (PCoA) plots using the Bray-Curtis distance, comparing with the uncorrected and ConQuR-corrected datasets. Assessed by the PERMANOVA R^2^ (which evaluates the multivariate association between the microbial community and a certain variable, here quantifies the proportion of microbiome data variability explained by the variable), correction with TG leads to the most significant reduction in batch effects, which was expected. Visually, it not only equalizes the mean and dispersion (centroid and size of the ellipses) across groups but also removes higher-order batch/group effects (shown by angles of the ellipses). Correction for QuanT, SVA, and RUV’s estimated surrogate variables leads to comparable efficacy in reducing batch effects and maintaining the effects of high blood pressure (HBP).

It is worth noting that the CARDIA study has well-balanced batch designs; thus the analysis with NC leads to reasonable and comparable results as the analyses with adjustments.

### Application to HIVRC, an integrated dataset with multiple HIV studies

Our second example used data from the HIV re-analysis consortium (HIVRC)^5^. In this consortium study, raw 16s rRNA gene sequencing data were solicited from 22 studies that investigated the relationship between the gut microbiome and HIV infection. Data from 17 of these studies were subsequently processed through a common pipeline with taxa assigned using Resphera Insight^35^. Note that the studies in the final-processed data have distinct experimental designs and sequencing protocols (Supp. Tab. 2 of Tuddenham, et al. 2020^17^). Thus, the data have more pronounced variability than that observed within a single study, such as the CARDIA study.

We selected two studies with similar library sizes (5000-5500) but very different study designs (an observational study in the US^36^ and a clinical trial in Europe^37^), which include 37 and 24 samples respectively. After preprocessing, the data contains 126 taxa. We applied QuanT, SVA, and RUV to the data aggregated at the genus level. HIV status (Negative = 0, Positive = 1) was regarded as the variable of interest, while age, gender and BMI were considered as covariates. We retained complete cases only and conducted the same set of analyses as on the CARDIA data.

Fig. 3a and 3b compare the consistency of differential abundance analysis results obtained analyzing these data using various methods. Fig. 3a shows the number of differentially abundant taxa identified in the HIVRC dataset by Benjamini-Hochberg adjusted p-value < 0.05. We identified 19 discoveries adjusting for true group indicators (TG), which were considered as the gold standard. QuanT leads to 17 true discoveries and 1 false discovery. NC leads to 11 true discoveries and 16 false discoveries. SVA leads to 11 true discoveries and 21 false discoveries. RUV leads to 8 true discoveries and 5 false discoveries. Therefore, QuanT again performs better than the other methods when applied to the HIVRC dataset and has the lowest FDR compared to competitors. In Fig. 3b, we consider the top K detected differentially abundant taxa. QuanT is the most consistent with the gold standard, TG, followed by NC, SVA, and RUV.

**Fig. 3.**
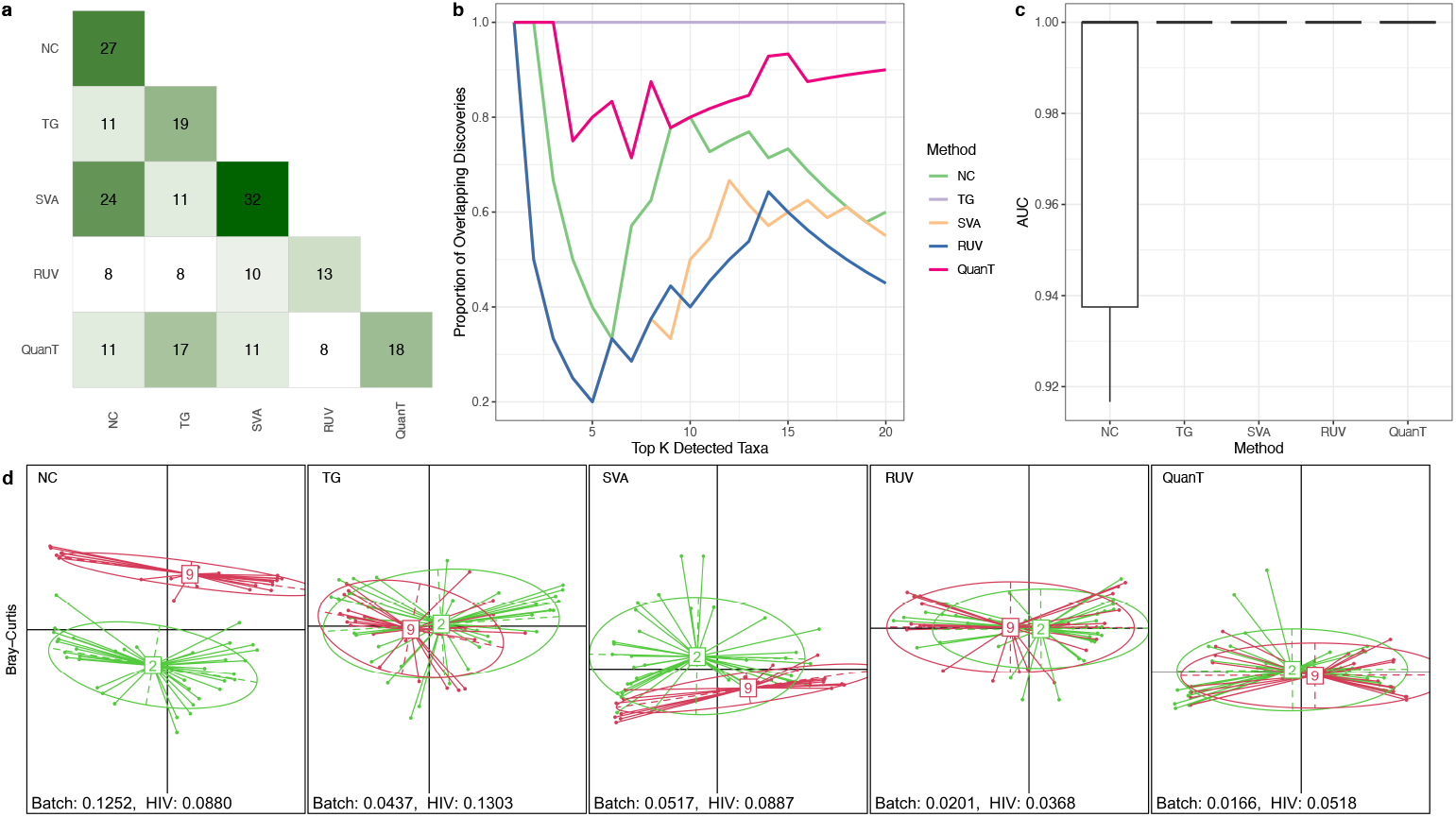
Evaluation on the HIVRC data. NC: no batch correction; TG: correction using true group indicators. **a** Heatmap representing the number of differentially abundant taxa identified in the HIVRC dataset by Benjamini-Hochberg adjusted p-value < 0.05; diagonal elements are numbers detected, off-diagonal elements are numbers detected in common. **b** Proportions of overlapping taxa in the top K detected differentially abundant taxa between each method and TG in the HIVRC dataset. **c** Average cross-validated area under the receiver operating characteristic curves (ROC-AUC) of predicting HIV from the ConQuR-corrected data via random forest. Larger values indicate stronger predictive capabilities. **d** PCoA plots of the ConQuR-corrected data based on the Bray-Curtis dissimilarity, with PERMANOVA R^2^ of study ID and HIV.

Fig. 3c shows the average cross-validated ROC-AUC of predicting HIV from the ConQuR-corrected data via random forest. TG, SVA, RUV, and QuanT all perform perfectly and are better than no correction. Fig. 3d shows the PCoA plots using Bray-Curtis distance, comparing with the uncorrected and ConQuR-corrected datasets. Visually, TG, RUV and QuanT lead to better harmonization of the microbiome data from the two distinct HIV studies. As quantified by PERMANOVA R^2^, QuanT leads to the greatest reduction of the effects of the unwanted heterogeneity, followed by RUV, SVA and TG. On the other hand, SVA maintains the HIV effects the most, followed by TG, QuanT and RUV (figure 3d).

### Application to an integrated microbiome dataset in colon cancers

Our third example came from two colorectal cancer (CRC) studies^38, 39^ from the curatedMetagenomicData package^40^. The CuratedMetagenomicData data package offers standardized, curated, and analysis-ready metagenomic datasets derived from human microbiome studies, facilitating comprehensive meta-analyses and cross-study comparisons by ensuring consistent data processing and annotation. The bacterial, fungal, and archaeal taxonomic abundances for each sample were calculated with MetaPhlAn3^41^, and metabolic functional potential was calculated with HUMAnN3^41^. We selected two CRC studies^38, 39^ from this package, which include 154 and 80 stool samples, respectively. After filtering, 166 genera were included in the data. These studies collectively cover samples from three distinct health conditions: colorectal cancer (CRC), adenoma, and disease-free controls, with CRC subdivided into adenocarcinoma and carcinoma, and adenoma into simple adenoma and advanced adenoma.

We applied QuanT, SVA, and RUV to this CRC dataset to find the hidden heterogeneity directions, treating the true study indicators as unknown. We considered the five phenotype categories (adenocarcinoma, carcinoma, adenoma, advanced adenoma, control) as the primary variable of interest, age (Adult = 0, Senior = 1) and gender (Male = 0, Female = 1) as the covariates to adjust for.

Fig. 4a and 4b compare the consistency of differential abundance analysis results across various methods. Fig. 4a shows the number of differentially abundant taxa identified in the CRC dataset by Benjamini-Hochberg adjusted p-value < 0.05. TG (gold standard) leads to 8 discoveries. QuanT leads to 5 true discoveries with 1 false discovery. NC leads to 6 true discoveries and 24 false discoveries. SVA leads to 6 true discoveries and 23 false discoveries. RUV leads to 1 true discovery with no false discovery. Therefore, NC and SVA lead to inflated false discovery proportion when applied to the CRC dataset. RUV and QuanT have reasonable false discovery proportion control, while QuanT is notably better at identifying true signals than RUV. In Fig. 3b, we consider the top K detected differentially abundant taxa. QuanT is the most consistent with the gold standard, TG. NC, SVA, and RUV perform similarly are all outperformed by QuanT.

**Fig. 4.**
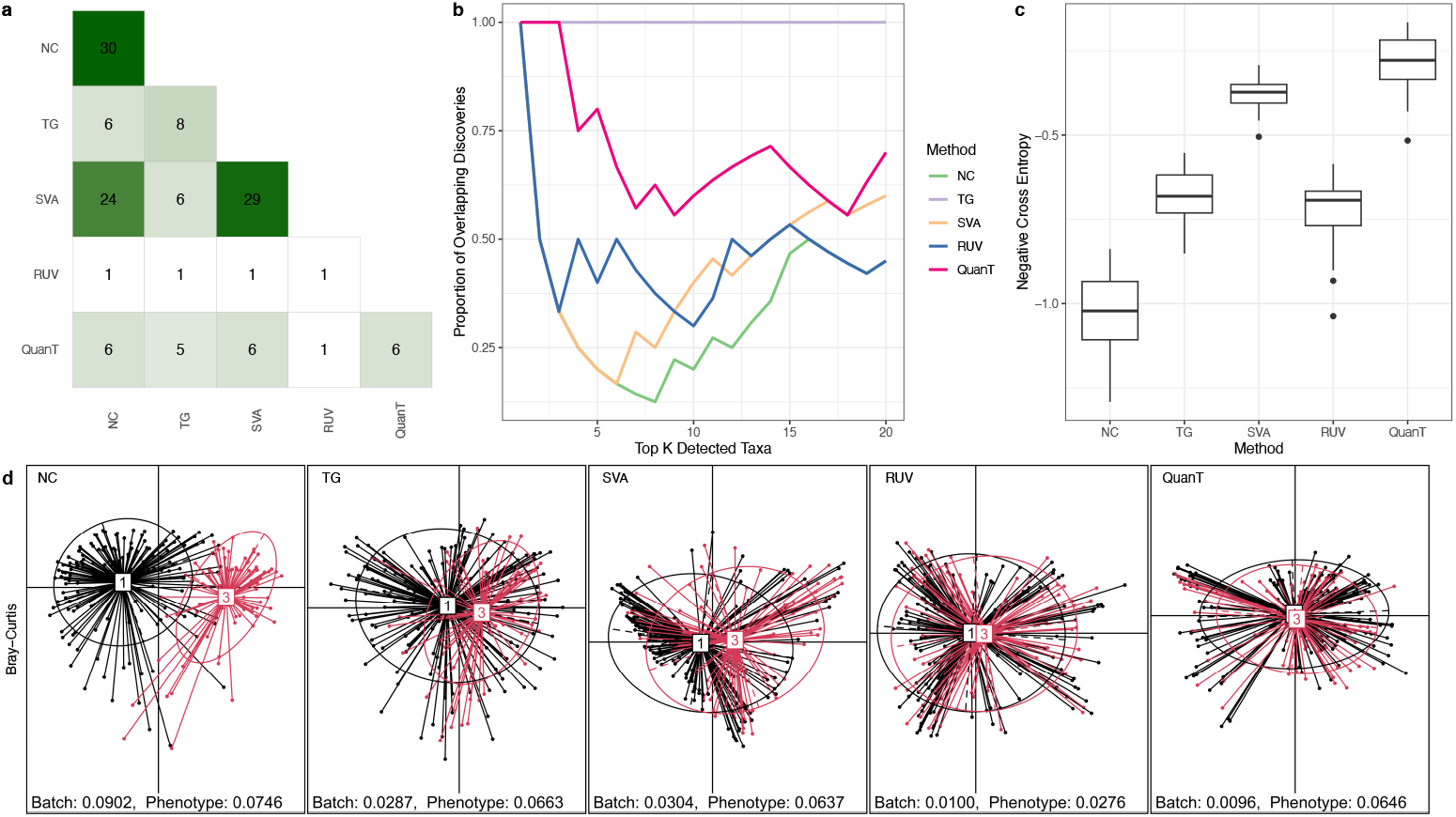
Evaluation on the CRC data. NC: no batch correction; TG: correction using true group indicators. **a** Heatmap representing the number of differentially abundant taxa identified in the CRC dataset by Benjamini-Hochberg adjusted p-value < 0.05; diagonal elements are numbers detected, off-diagonal elements are numbers detected in common. **b** Proportions of overlapping taxa in the top K detected differentially abundant taxa between each method and TG in the CRC dataset. **c** Average cross-validated negative cross entropy of predicting HIV from the ConQuR-corrected data via random forest. Larger values indicate stronger predictive capabilities. **d**. PCoA plots of the ConQuR-corrected data based on the Bray-Curtis dissimilarity, with PERMANOVA R^2^ of study ID and Phenotype.

Fig. 4c presents the prediction accuracy, measured by negative cross entropy, for the five phenotype groups using the original and ConQuR-corrected CRC microbiome data. Here, QuanT performs the best, followed by SVA, RUV, TG, and NC. Fig. 4d displays the PCoA plots using Bray-Curtis distance. Visually, QuanT leads to similar data harmonization as using the true study IDs, and outperforms SVA, RUV and NC. This is numerically confirmed by the PERMANOVA R^2^ of the study ID and the phenotypes of interest.

### Evaluation using simulated data

We applied SVA, RUV and QuanT to the simulated datasets to identify the underlying hidden heterogeneity, treating the batch variables as unknown. Like in the CARDIA, HIVRC, and CRC analyses, we used linear regression models to conduct differential abundance analysis, adjusting for hidden variables estimated by SVA, RUV, and QuanT. When data are simulated from MIDASim, as shown in Fig. 5a, QuanT leads to better FDR control compared to SVA and RUV. Fig. 5b shows that the discoveries by QuanT align more closely with the gold standard than SVA, RUV, and the NC methods. Here, the gold standard varies: in the absence of batch effects, NC is the gold standard, while when batch effects are present, the model correcting for true batch ID is the gold standard. More simulation results based on MIDASim (Supp. Fig. 2,4,6) and results based on parametric simulations (Supp. Fig. 1,3,5) lead to similar conclusions.

**Fig. 5.**
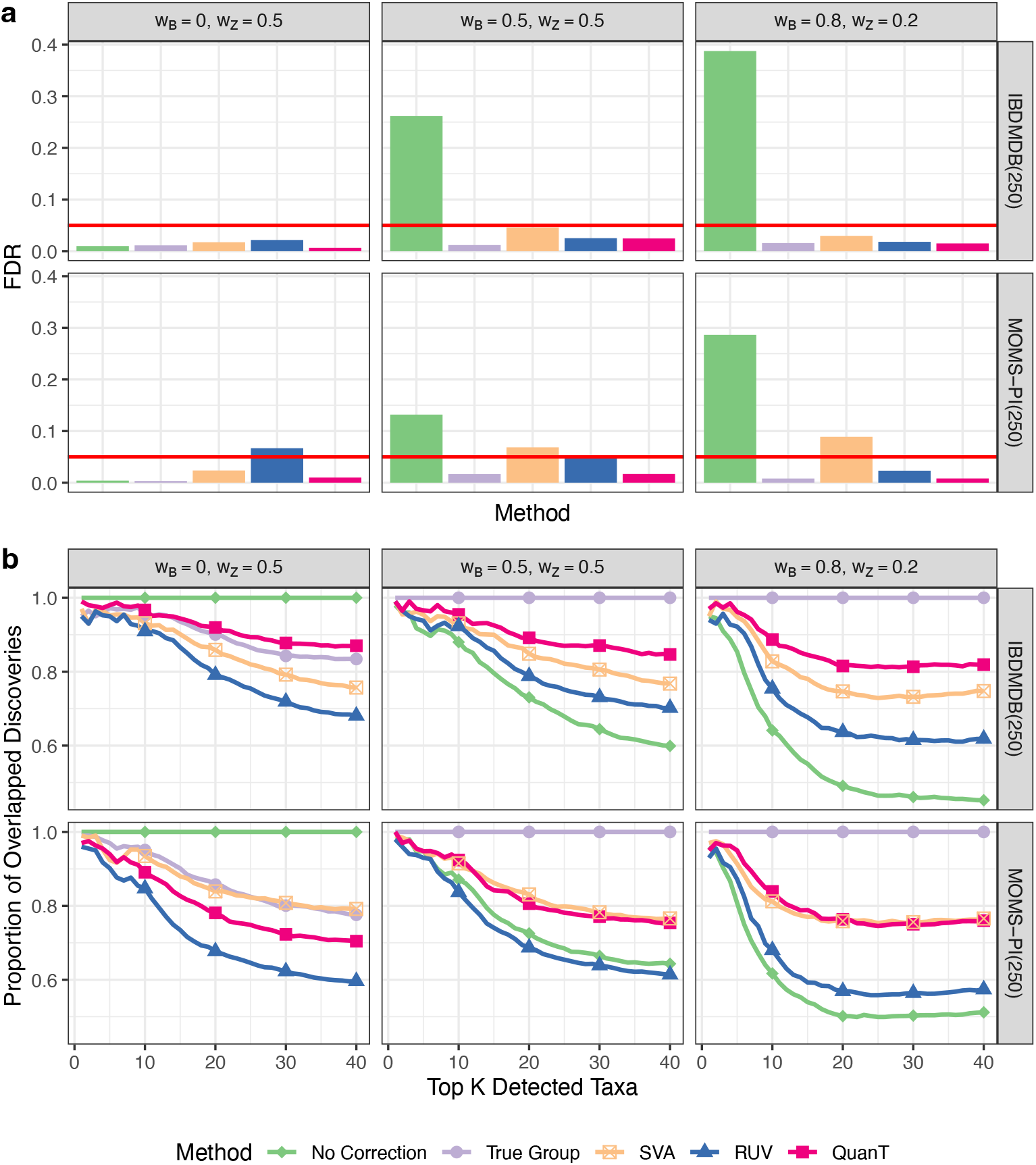
Evaluation on the simulated data. The data are simulated by MIDASim using the IBDMDB and MOMS-PI datasets as templates. **a** FDR of the differential abundance analysis via adjusted linear regression with BH-adjusted p-value < 0.05. **b** Proportions of overlapping taxa in the top K detected differentially abundant taxa between each method and the gold standard in simulated data. Simulation setups vary across different datasets (rows), and different combinations of batch and key variable effects (columns).

We also applied ConQuR to remove the true batch, SVs and QSV effects and generate a “batch-free” dataset. Integrating QuanT with ConQuR in simulations (Supp. Fig. 7-10) yielded microbial profiles where confounding batch effects were eliminated, while the effects of the variable of interest were preserved. Finally, QuanT demonstrates scalable computational performance for large-scale microbiome studies, as evidenced in Tab. 1 and Tab. 2.

For our simulation scenario 3, we applied LinDA^42^ and LDM-CLR^43^, two modern tools that are designed for compositional hypothesis in microbiome data. These results (Supp. Fig. 11-16) align with the conclusions from our other simulation studies.

In short, QuanT demonstrates improved performance compared to existing methods, on simulated datasets and biological datasets, regardless of the effect size and the type of source of unmeasured heterogeneity.

### Computation of QuanT

We have streamlined the process of quantile regression and matrix decomposition within QuanT by employing strategies for enhanced efficiency. The algorithm executes quantile regression across various taxon and quantile levels independently, which lends itself well to parallel processing. We leverage the doParallel package in R to expedite this computation. In the second stage of QuanT, where SVD is performed, we focus on a few leading singular vectors because it is generally improbable for a singular vector that accounts for minimal variance to represent a critical direction established in the first stage. Consequently, it is sufficient to compute only the leading singular vectors of the binarized residual matrix during the correlation calibration phase. For this reason, QuanT only calculates the top 15 principal components using the RSpectra R package.

Tables 1 and 2 present the computational time of QuanT for datasets with different numbers of samples and taxa. Table 1 illustrates the scalability of QuanT’s computational time with respect to various numbers of samples (*n*) and taxa (*p*), benchmarked against SVA and RUV. For each configuration of *n* and *p*, we generated ten datasets using the DM simulation with parameters (*w*_*B*_, *w*_*z*_) = (0.5,0.25) and the IBDMDB dataset as the template. We also compared the computational time of QuanT, SVA, and RUV for analyzing the three real-world datasets (CARDIA, HIVRC, and CRC) (Tab. 2). Overall, QuanT exhibits slower computation than SVA and RUV due to its multiple quantile regression procedures. Nevertheless, the run time remains practical and the method is applicable to reasonably large microbiome datasets.

**Table 1.**
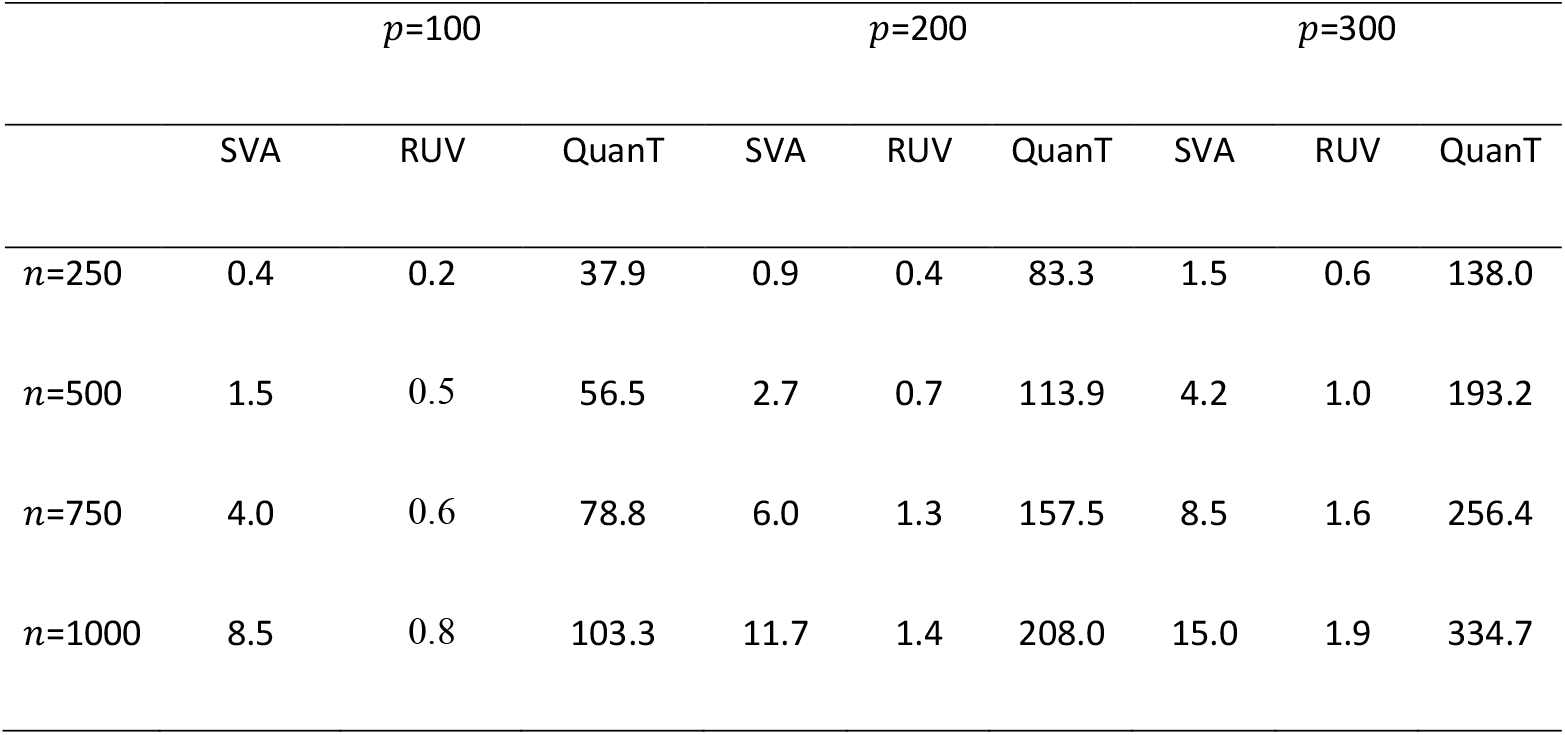
The computation time (in seconds) of QuanT with different numbers of samples and taxa, compared with SVA and RUV. The data are simulated from the Dirichlet-Multinomial distribution using the IBDMDB dataset as the template.

**Table 2.**
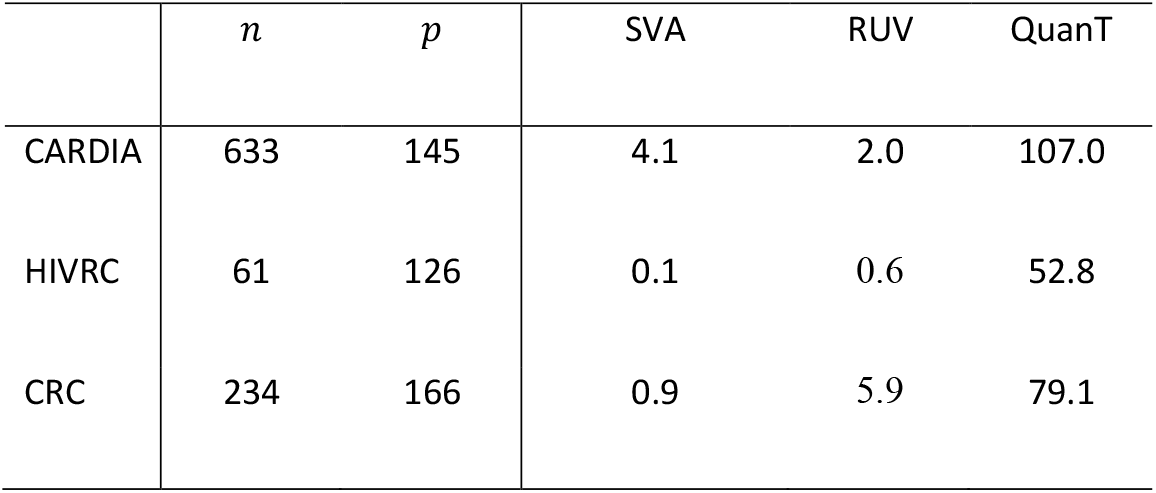
The computation time (in seconds) of QuanT for the three datasets analyzed, compared with SVA and RUV.

## Discussion

Identifying unmeasured heterogeneity is a challenging yet important task in microbiome data analysis. Failing to identify and adjust for the unmeasured heterogeneity can drastically increase the false discovery rate and hinder the detection of true signals. Due to lack of methods specifically tailored to microbiome data, researchers often use methods developed for genomic studies. However, these methods do not account for the challenging characteristics of microbiome data, such as sparsity and overdispersion. In addition, they focus solely on the information underlying the mean expression and ignore other aspects of the data distribution. As shown the “Results” section, these methods lead to increased false discoveries and/or reduced power when analyzing microbiome studies.

We present QuanT, the first comprehensive tool to identify unmeasured heterogeneity in microbiome data. QuanT uses quantile regression models and so can thoroughly investigate the underlying data structure at multiple quantiles. QuanT outputs a set of vectors as surrogates (QSV) for the unmeasured heterogeneity, which offers users the flexibility to incorporate them in different types of downstream analysis. These vectors can be treated as known covariates and adjusted for in regression models. They can also be used in conjunction with a batch effect removal method, such as ConQuR, to generate microbial profiles without the effects of hidden heterogeneity.

We applied QuanT to three microbiome datasets and a wide range of simulated datasets to assess its performance, in comparison with competing methods. The microbiome datasets include large-scale epidemiological studies with samples processed in different sequencing runs, and curated microbiome datasets integrated from various publicly available studies. Our simulations are comprehensive, generating microbiome profiles that mimic real-world microbiome studies in gut and vaginal environments and using two distinct simulation mechanisms. Visually and numerically, we showed that QuanT improved subsequent differential abundance analysis, offering better FDR control and improved statistical power, and providing results that align better with the gold standard in most setups. Frequently, the results of differential abundance analysis following QuanT are comparable to those when the true labels of batches or studies are known. It also offers comparable or improved performance in predicting the variable of interest, as well as reducing the impact of hidden heterogeneity while preserving key signals at the community level.

Despite its advantages, QuanT has several limitations and potentials that could be investigated in future research. First, nowadays, it is common that microbiome studies are conducted in a longitudinal manner to investigate the dynamics of the microbiome. Such studies typically produce abundance data collected across multiple time points from the same individuals. For longitudinal data, one tentative approach is to incorporate time as an additional adjustment variable into the QuanT framework. However, this approach ignores the correlations between repeated samples from the same person and may lead to spurious conclusions. In contrast, in a well-designed study, multiple samples from the same individual are usually sequenced on the same plate. In such cases, baseline microbiome data can be used for the identification of unmeasured heterogeneity. Sensitivity analyses may then be conducted by leveraging microbiome samples that share similar batch structures—for example, by performing analyses separately on baseline samples and on samples collected at later time points, thereby assessing the consistency of results across these subsets. Secondly, QuanT leads to well-controlled false discovery rate in our real and extensive simulation experiments, although it lacks theoretical guarantee. However, theoretical guarantees might be derived given strict assumptions about the data generation mechanism, which, in practice, is usually less important. Finally, a potential extension of QuanT is to identify unmeasured heterogeneity in gene expression data, especially in single cell and spatial transcriptomics data which have similar sparsity and over-dispersion as do microbiome data. It would be worth investigating how QuanT performs when applied to these data types, which might require some adaptations. However, these aspects are beyond the scope of this paper, and we leave them to future studies.

## Conclusions

In summary, we propose a versatile approach, QuanT, to uncover concealed heterogeneities within microbiome data. QuanT stands out from existing approaches by detecting hidden data structures beyond the mean and by easing the constraints of distributional assumptions. When integrated into the analysis pipeline, QuanT effectively mitigates the confounding influences of undesired latent variables, enhances false discovery rate control, and preserves statistical power.

## List of abbreviations

QuanT: quantile thresholding
SVA: surrogate variable analysis
RUV: remove unwanted variation
SVD: singular value decomposition
iSV: iterative singular vectors
QSV: quantile surrogate variables
VOI: variable of interest
CARDIA: Coronary Artery Risk Development in Young Adults
CVD: cardiovascular disease
HBP: high blood pressure
TG: true group
NC: no correction
ROC-AUC: area under the receiver operating characteristic curve
PCoA: principal coordinate analysis
HIVRC: HIV re-analysis consortium
CRC: colorectal cancer
IBDMDB: Inflammatory Bowel Disease Multi-omics Database
MOMS-PI: Multi-Omic Microbiome Study: Pregnancy Initiative
DM: Dirichlet-Multinomial
MIDASim: Microbiome Data Simulator

## Declarations

### Ethics approval and consent to participate

Not applicable. Because this study only involved secondary analysis of existing, de-identified datasets, it is not considered human subject research.

### Consent for publication

Not applicable.

### Availability of data and material

The CARDIA data can be shared upon request by submitting a CARDIA Data Set Request-Intent to Analyze Form to the CARDIA Coordinating Center, University of Alabama at Birmingham, at https://www.cardia.dopm.uab.edu/publications-2/publications-documents.

The HIVRC data are available in Synapse under accession code syn18406854. The CRC data are from the curatedMetagenomicData package at https://waldronlab.io/curatedMetagenomicData/. The Inflammatory Bowel Disease Multiomics Database (IBDMDB) project^28^ and the vaginal microbiome dataset from Multi-Omic Microbiome Study^29^: Pregnancy Initiative (MOMS-PI) project, which were used as templates for simulation studies, are provided in the Bioconductor R package, HMP2Data^44^.

The R package QuanT is available at https://github.com/jiuyaolu/QuanT in formats appropriate for Macintosh, Windows, or Linux systems. A vignette demonstrating use of the package is included (can be accessed at https://jiuyaolu.github.io/QuanT/QuanT.html).

### Competing interests

The authors declare that they have no competing interests.

### Funding

This work was supported in part by R01GM147162 (Lu, Satten, and Zhao), R01HL155417 (Ling), R01GM151301 (Ling) and R01GM155734 (Ling). The Coronary Artery Risk Development in Young Adults Study (CARDIA) is supported by contracts 75N92023D00002, 75N92023D00003, 75N92023D00004, 75N92023D00005, and 75N92023D00006 from the National Heart, Lung, and Blood Institute (NHLBI). The HIVRC data used in this study are from work that was supported by the HIV Microbiome Re-analysis Consortium.

### Authors’ contributions

JL, GS, NZ and WL developed the method, analyzed the data, and wrote the manuscript. JL prepared the software package and vignette. LL and KM contributed to the dataset generation and manuscript development with technical input. All authors read and approved the final manuscript.

## Acknowledgements

The authors thank Drs. Susan A. Tuddenham, James White, Wei Li A Koay, Cynthia L. Sears, and all members on the HIV Microbiome Re-analysis Consortium for collecting and processing the HIVRC dataset.

## Notes

### Competing Interest Statement

The authors have declared no competing interest.

### Summary of Updates

Updated the algorithm and the results.

