## Supplementary Material for "Identifying unmeasured heterogeneity in microbiome data via quantile thresholding (QuanT)"

### Simulation results

#### Results based on regression

Supp. Fig. 1 and Supp. Fig. 2 show the false discovery rate (FDR) from downstream differential abundance analysis adjusted for estimated hidden heterogeneity by SVA, RUV, and QuanT, based on simulations using Dirichlet-Multinomial distribution and MIDASim, respectively.

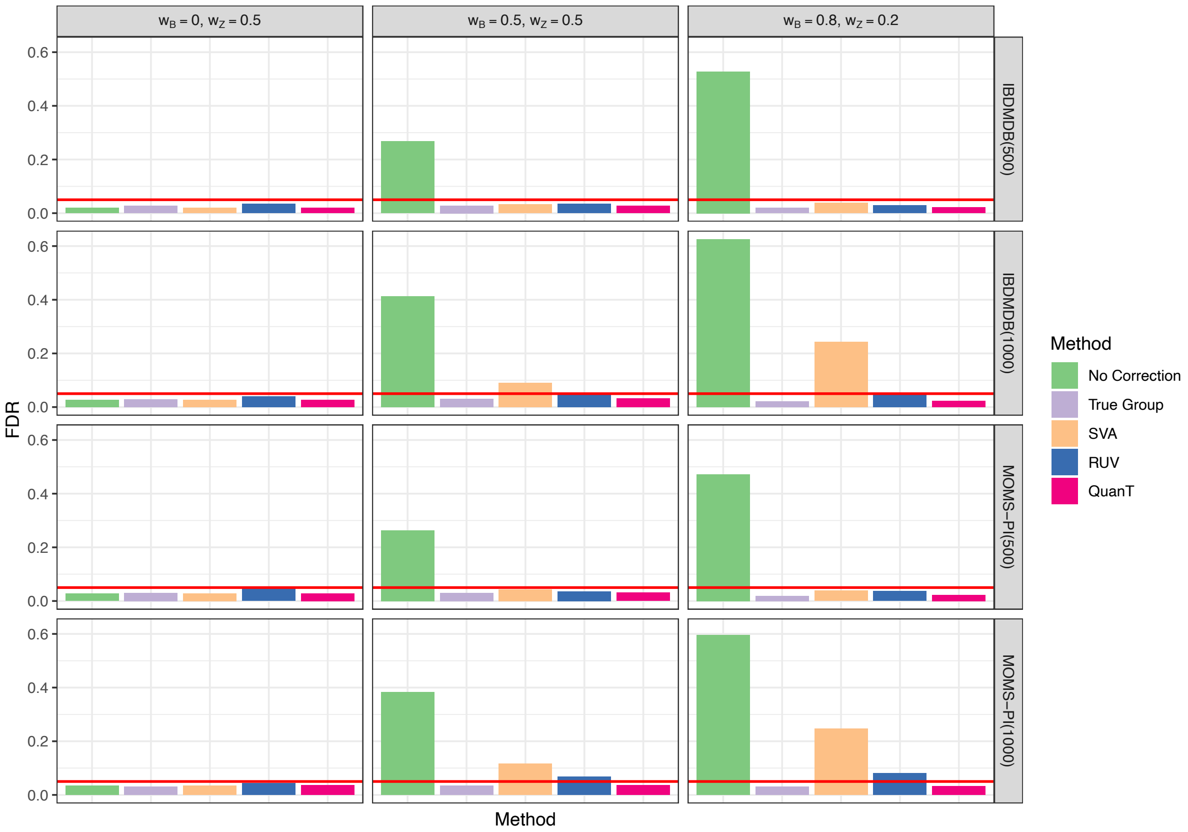


**Supp. Fig. 1 | FDR of downstream differential abundance analysis adjusted for different estimated hidden heterogeneity when data are simulated from the Dirichlet-Multinomial**

**distribution using the IBDMDB and MOMS-PI datasets as the template. $w_B$ and $w_z$ define the underlying batch effect and clinical effect. The number in paratheses is the sample size used in the simulations.**

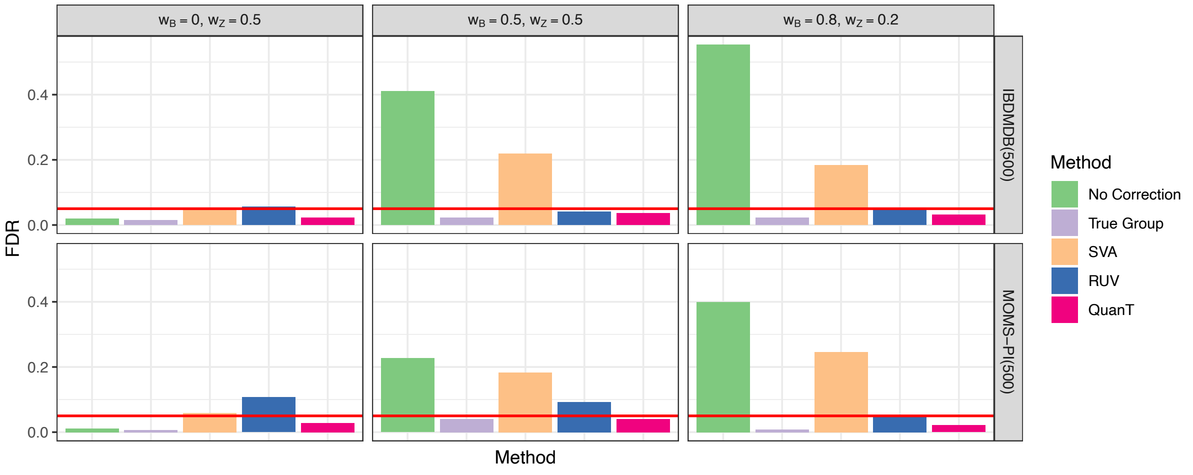


**Supp. Fig. 2 | FDR of downstream differential abundance analyses adjusted for different estimated hidden heterogeneity when data are simulated by MIDASim using the IBDMDB and MOMS-PI datasets as the template. $w_B$ and $w_z$ define the underlying batch effect and clinical effect. The number inside paratheses is the sample size use in the simulations.**

Supp. Fig. 3 and Supp. Fig. 4 show the sensitivity of downstream differential abundance analyses using data generated with the Dirichlet-Multinomial distribution and MIDASim, respectively. Since no correction, SVA, and RUV do not control FDR, they are omitted from the comparison. The figures illustrate that adjustment using QuanT's surrogate variables leads to power that is comparable to adjusting for the true batch variable.

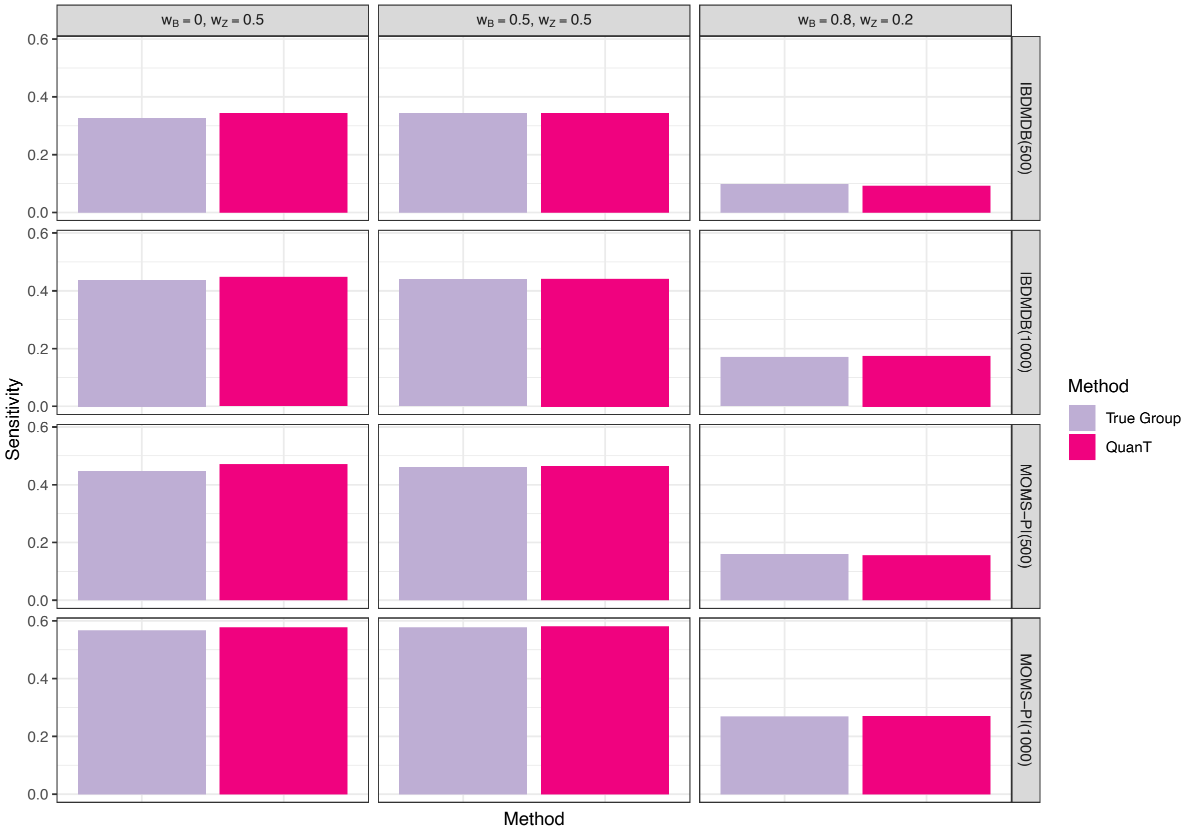


**Supp. Fig. 3 | Sensitivity of downstream differential abundance analyses adjusted for different estimated hidden heterogeneity when data are simulated from the Dirichlet-Multinomial distribution using the IBDMDB and MOMS-PI datasets as the template. $w_B$ and $w_z$ define the underlying batch effect and clinical effect. The number in paratheses is the sample size used in the simulations.**

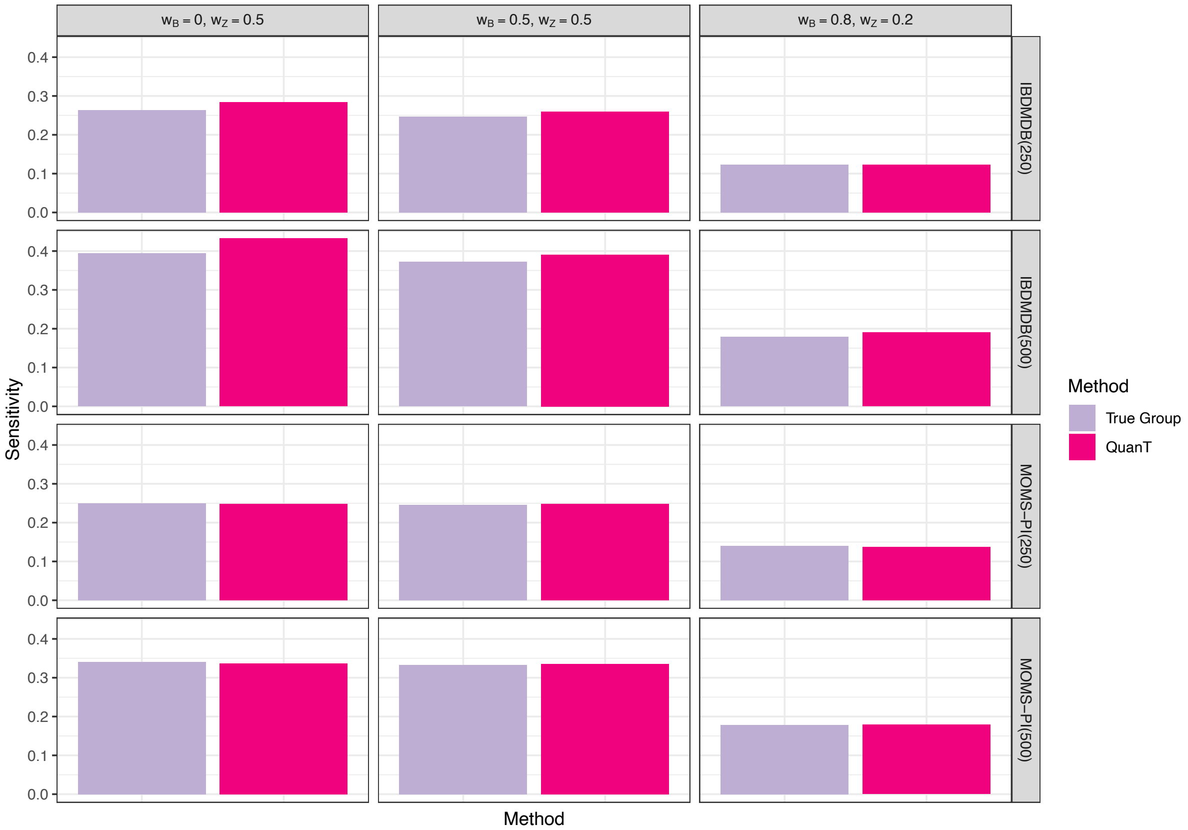


**Supp. Fig. 4 | Sensitivity of downstream differential abundance analyses adjusted for different estimated hidden heterogeneity when data are simulated by MIDASim using the IBDMDB and MOMS-PI datasets as the template. $w_B$ and $w_z$ define the underlying batch effect and clinical effect. The number in paratheses is the sample size in simulations.**

Supp. Fig. 5 and Supp. Fig. 6 show the proportions of overlapping discoveries among the top K differentially abundant taxa identified by each method, compared with the gold standard, when data were generated using the Dirichlet-Multinomial distribution and MIDASim. The corresponding results for Dirichlet-Multinomial simulations are shown in Fig. 2c. When there are no batch effects and the no correction method is the gold standard, SVA and QuanT performance well, followed by adjustment using a "true" batch variable. RUV leads to the least consistent results with the gold standard when there are no batch effects. Note that in such a situation, the so-called “true” batch is just a binary indicator that is simulated with no impact on the microbiome data but is associated with the variable of interest. When there are actual batch effects present and correcting for the true batch variable is the gold standard, QuanT produces a list of differentially abundant taxa that agrees best with the gold standard, followed by SVA and RUV.  No correction yields the worst results when there are batch effects.

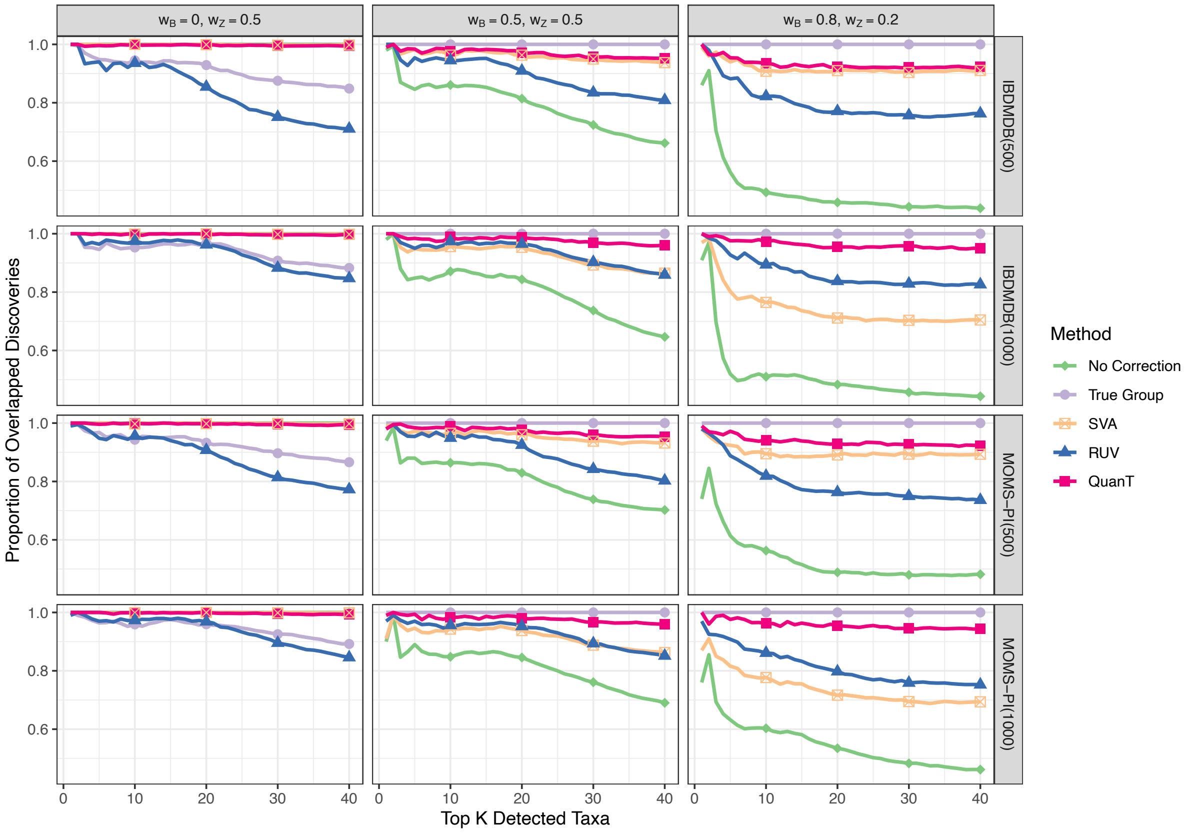


**Supp. Fig. 5 | Proportions of overlapping discoveries among the top K differentially abundant taxa identified by each method, compared with the gold standard when data are simulated from the Dirichlet-Multinomial distribution using the IBDMDB and MOMS-PI**

**datasets as the template. $w_B$ and $w_z$ define the underlying batch effect and clinical effect. When there is no batch effect ($w_B = 0$), the method with no correction is the gold standard. When batch effect exists ($w_B \neq 0$), the method with true batch correction is the gold standard.**

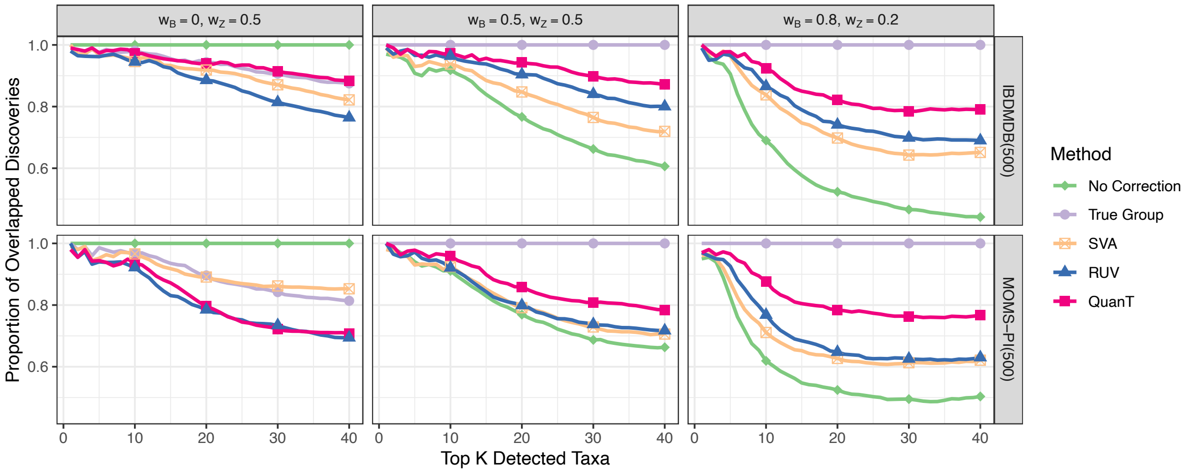


**Supp. Fig. 6 | Proportions of overlapping discoveries among the top K differentially abundant taxa identified by each method, compared with the gold standard when data are simulated by MIDASim using the IBDMDB and MOMS-PI datasets as the template. $w_B$ and $w_z$ define the underlying batch effect and clinical effect. When there is no batch effect ($w_B = 0$), the method with no correction is the gold standard. When batch effect exists ($w_B \neq 0$), the method with true batch correction is the gold standard.**

**Results based on ConQuR correction**

Supp. Fig. 7 and Supp. Fig. 8 show violin plots for the PERMANOVA $R^2$ of the batch effects using the ConQuR-corrected relative abundance matrices from the Dirichlet-Multinomial and

MIDASim simulations. When there are batch effects, correcting for the true batch variable leads to the smallest $R^2$ for batch and no correction leads to the largest $R^2$ for batch as expected. QuanT and RUV have comparable performance, showing smaller $R^2$ of batch compared to SVA, which means that QuanT and RUV are more capable of identifying the hidden variables, and the effects of the identified variables can be successfully removed by ConQuR.

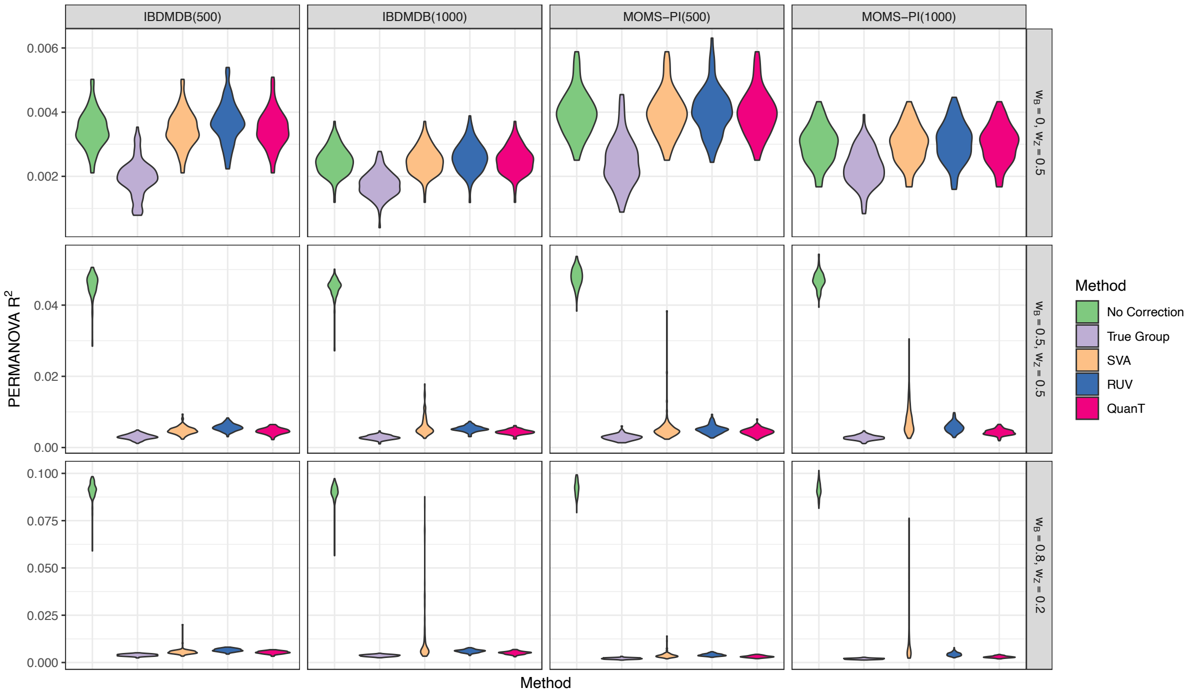


**Supp. Fig. 7 | PERMANOVA $R^2$ of the batch effects in the ConQuR-corrected data for the Dirichlet-Multinomial simulation using the IBDMDB and MOMS-PI datasets as the template. Here, ConQuR was used to remove the effects of hidden heterogeneity, detected via SVA, RUV, QuanT, or the true batch. When there is no correction, the original microbiome data was used for PERMANOVA. Parameters $w_B$ and $w_z$ define the underlying batch effect and clinical effect,. The violin plot was generated using 100 simulation replicates.**

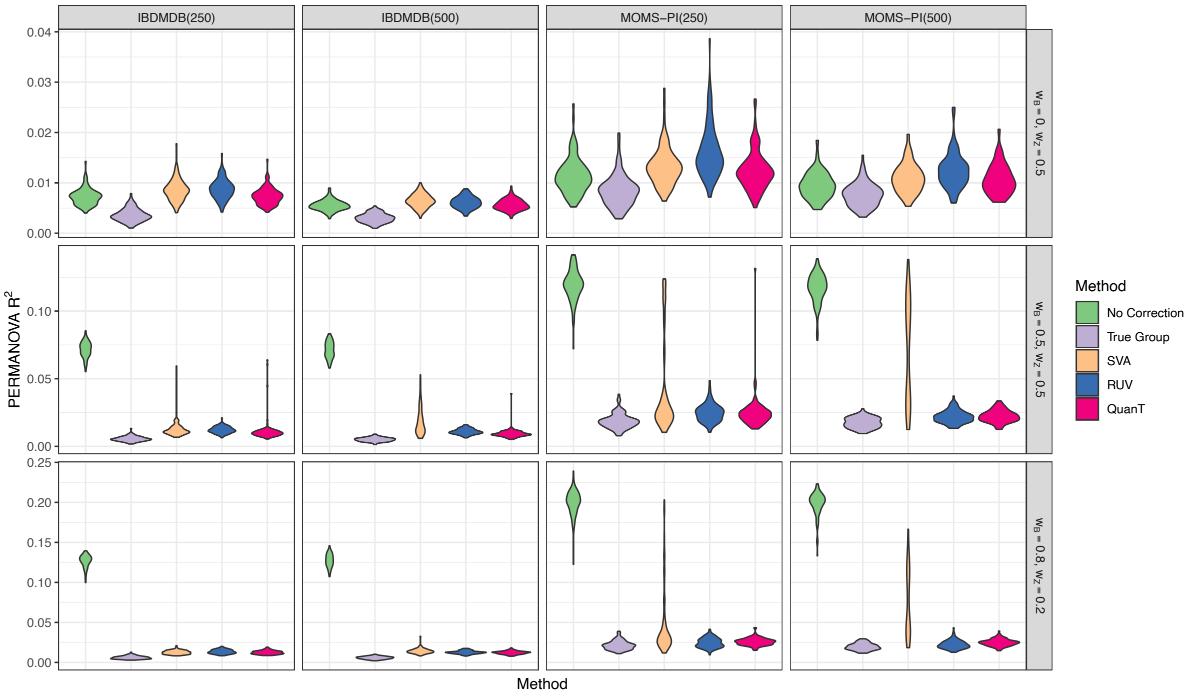


**Supp. Fig. 8 | PERMANOVA $R^2$ of the batch effects in ConQuR-corrected data when data are simulated by MIDASim using the IBDMDB and MOMS-PI datasets as the template. Here, ConQuR was used to remove the effects of hidden heterogeneity, detected via SVA, RUV, QuanT, or the true batch. When there is no correction, the original microbiome data was used for PERMANOVA. Parameters $w_B$ and $w_z$ define the underlying batch effect and clinical effect. The violin plot was generated using 100 simulation replicates.**

Supp. Fig. 9 and Supp. Fig. 10 show the PERMANOVA $R^2$ of the variable of interest in the ConQuR-corrected abundance matrices under the Dirichlet-Multinomial and MIDASim simulations. SVA, RUV, and QuanT all lead to similar $R^2$ of the variable of interest, which are comparable with correcting for the true batch variable.

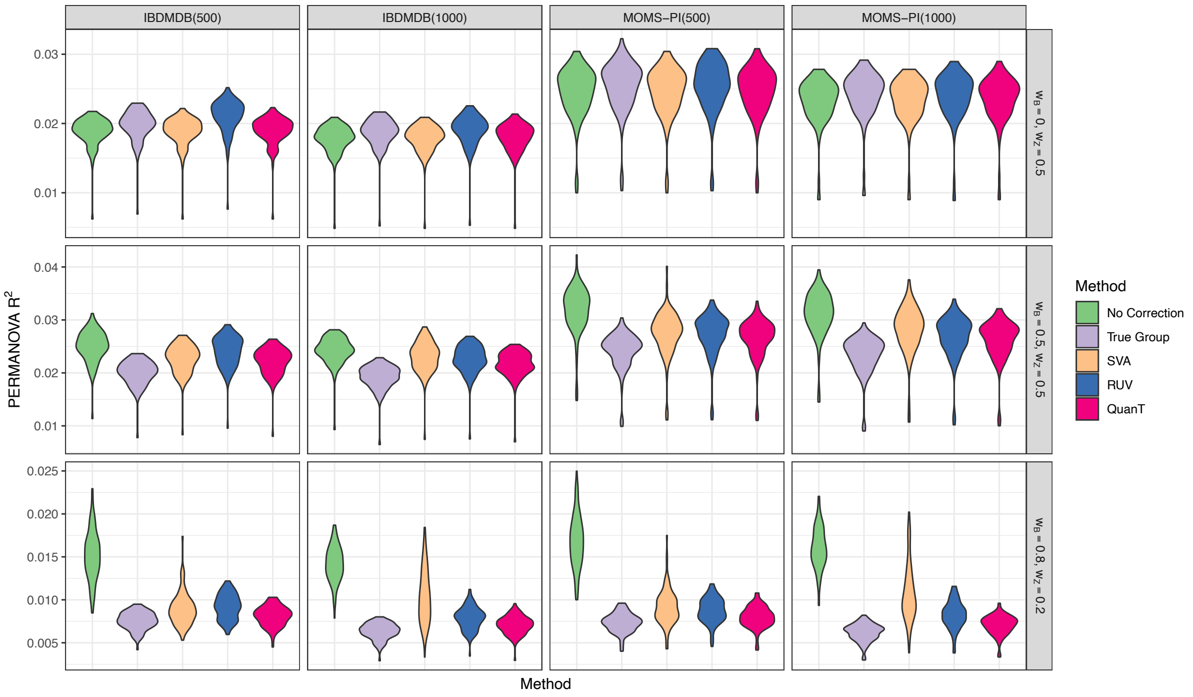


**Supp. Fig. 9 | PERMANOVA $R^2$ of the variable of interest in ConQuR-corrected data when data are simulated from the Dirichlet-Multinomial distribution using the IBDMDB and MOMS-PI datasets as the template. Here, ConQuR was used to remove the effects of hidden heterogeneity, detected via SVA, RUV, QuanT, or the true batch. When there is no correction, the original microbiome data was used for PERMANOVA. $w_B$ and $w_z$ define the underlying batch effect and clinical effect. The violin plot was generated using 100 simulation replicates.**

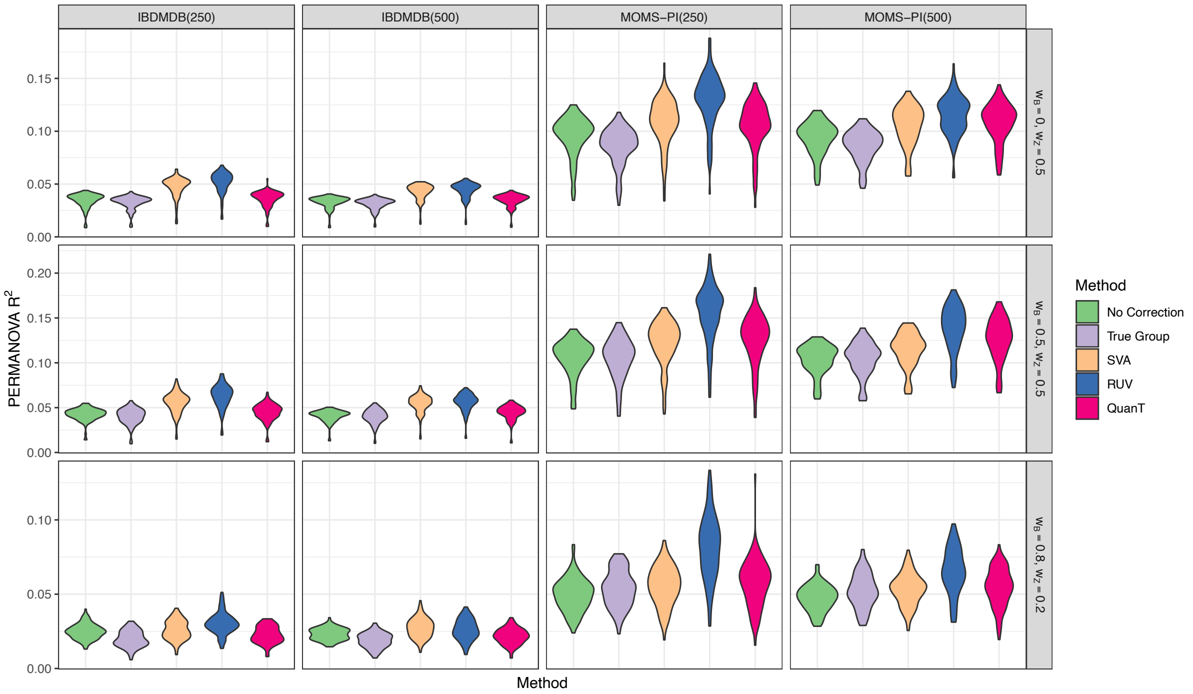


**Supp. Fig. 10 | PERMANOVA $R^2$ of the variable of interest in the ConQuR-corrected data when data are simulated by MIDASim using the IBDMDB and MOMS-PI datasets as the template. Here, ConQuR was used to remove the effects of hidden heterogeneity, detected via SVA, RUV, QuanT, or the true batch. When there is no correction, the original microbiome data was used for PERMANOVA. $w_B$ and $w_z$ define the underlying batch effect and clinical effect. The violin plot was generated using 100 simulation replicates.**

**Results based on compositional data simulation**

Same as the above simulations, we simulated data that mimic the two template datasets: IBDMDB and MOMS-PI. Data were simulated according to the third simulation scenario in the main manuscript, following Dirichlet Multinomial distribution. We subsequently applied SVA, RUV and QuanT to the simulated datasets to identify the underlying hidden heterogeneity, and used LinDA[1] and LDM-CLR[2] to conduct differential abundance analysis, instead of using linear regression models.

Supp. Fig. 11 and Supp. Fig. 12 show the false discovery rate (FDR) from downstream differential abundance analysis adjusted for estimated hidden heterogeneity by SVA, RUV, and QuanT, where differential abundance analysis is conducted with LinDA and LDM-CLR, respectively. Similarly to Supp. Fig. 1 and Supp. Fig. 2 , TG and QuanT have good FDR control, while the other methods fail to control the FDR in some settings.

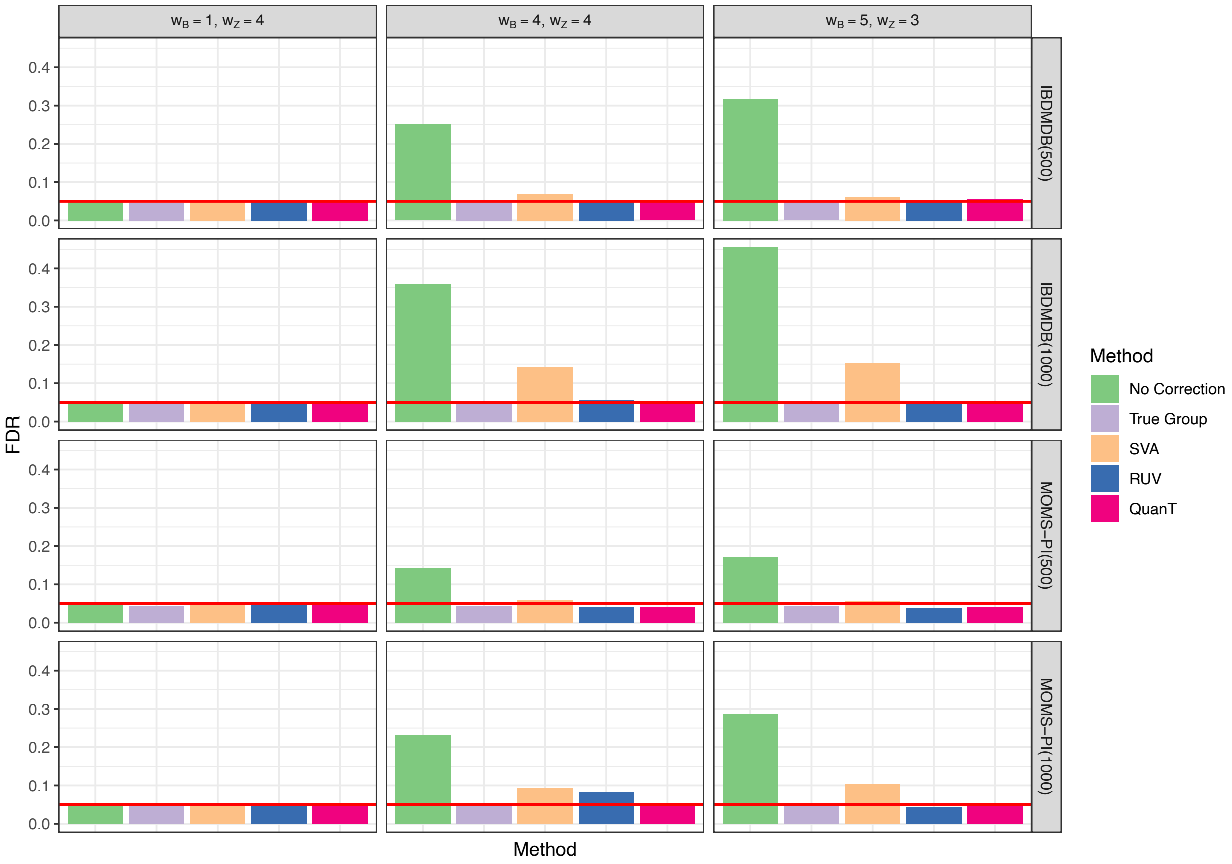


**Supp. Fig. 11 | FDR of downstream differential abundance analysis adjusted for different estimated hidden heterogeneity when data are simulated from the Dirichlet-Multinomial**

**distribution using the IBDMDB and MOMS-PI datasets as the template and analyzed with LinDA. $w_B$ and $w_z$ define the underlying batch effect and clinical effect. The number in paratheses is the sample size used in the simulations.**

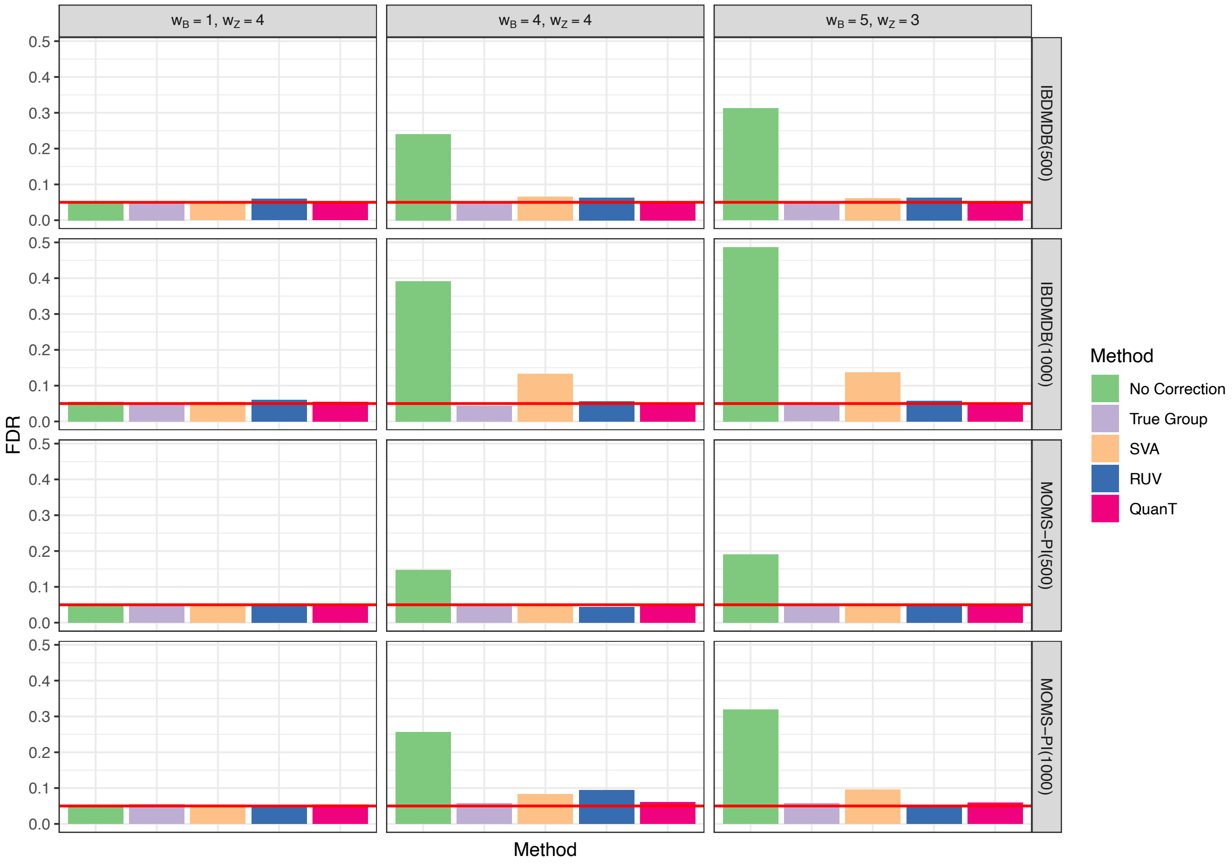


**Supp. Fig. 12 | FDR of downstream differential abundance analysis adjusted for different estimated hidden heterogeneity when data are simulated from the Dirichlet-Multinomial distribution using the IBDMDB and MOMS-PI datasets as the template and analyzed with LDM-CLR. $w_B$ and $w_z$ define the underlying batch effect and clinical effect. The number in paratheses is the sample size used in the simulations.**

Supp. Fig. 13 and Supp. Fig. 14 show the sensitivity of downstream differential abundance analyses where the analysis is conducted with LinDA and LDM-CLR, respectively. Since no correction, SVA, and RUV do not control FDR, they are omitted from the comparison. Similarly to Supp. Fig. 3 and Supp. Fig. 4, these figures illustrate that adjustment using QuanT's surrogate variables leads to power that is comparable to adjusting for the true batch variable.

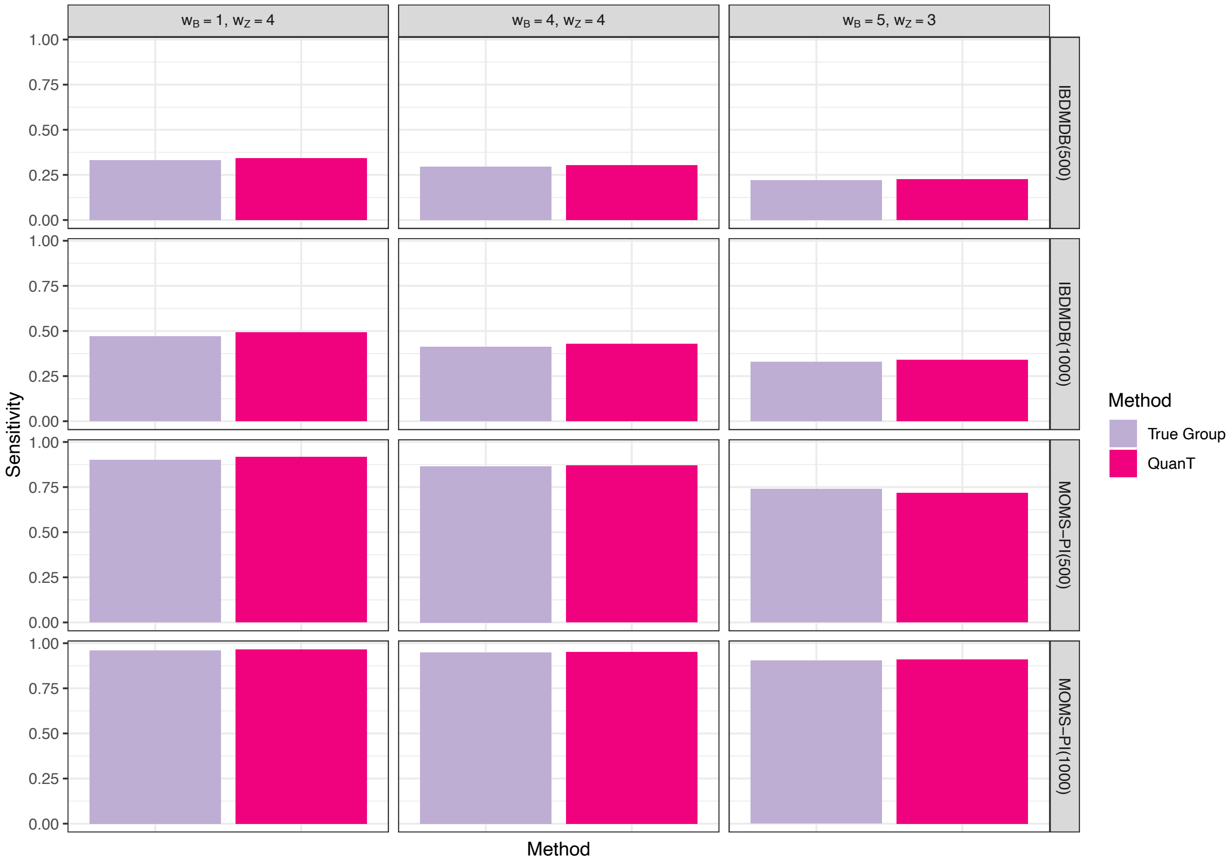


**Supp. Fig. 13 | Sensitivity of downstream differential abundance analyses adjusted for different estimated hidden heterogeneity when data are simulated from the Dirichlet-Multinomial distribution using the IBDMDB and MOMS-PI datasets as the template and analyzed with LinDA. $w_B$ and $w_z$ define the underlying batch effect and clinical effect. The number in paratheses is the sample size used in the simulations.**

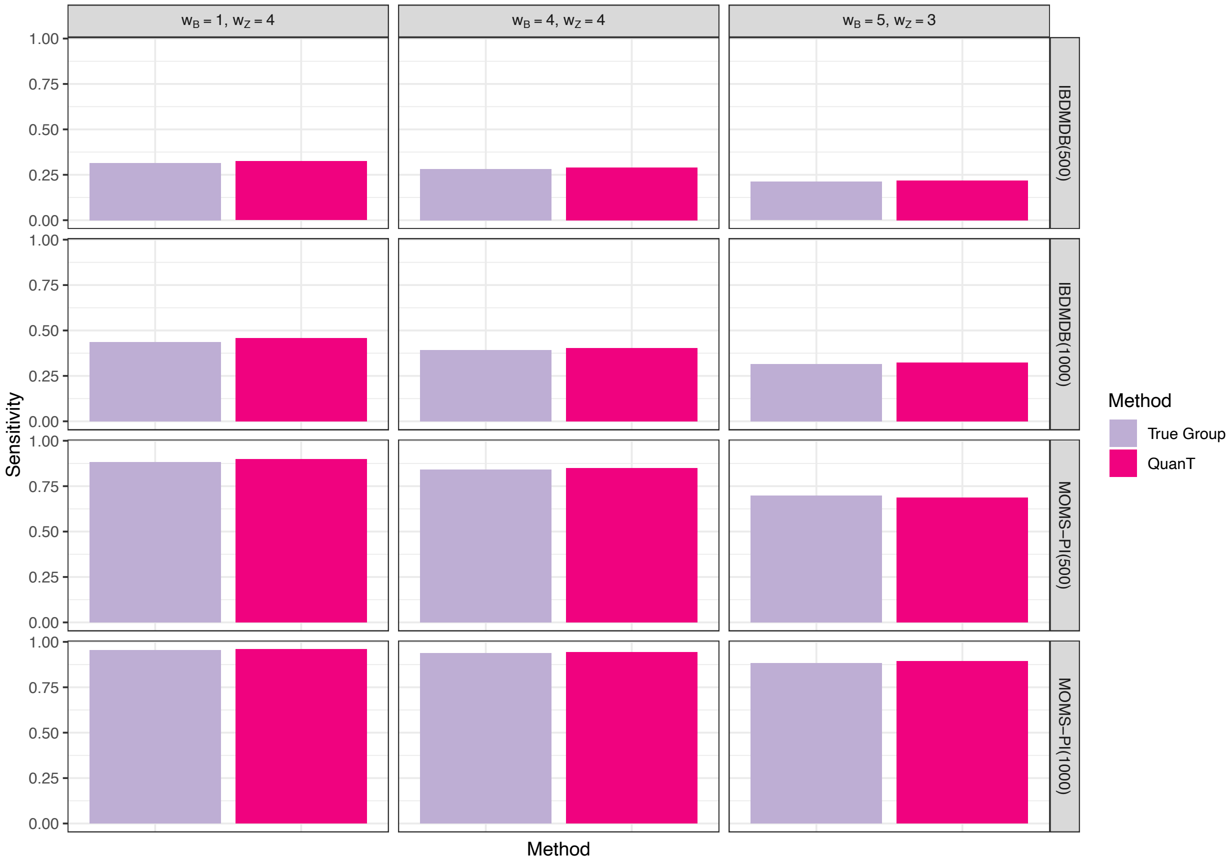


**Supp. Fig. 14 | Sensitivity of downstream differential abundance analyses adjusted for different estimated hidden heterogeneity when data are simulated from the Dirichlet-Multinomial distribution using the IBDMDB and MOMS-PI datasets as the template and analyzed with LDM-CLR. $w_B$ and $w_z$ define the underlying batch effect and clinical effect. The number in paratheses is the sample size used in the simulations.**

Supp. Fig. 15 and Supp. Fig. 16 show the proportions of overlapping discoveries among the top K differentially abundant taxa identified by each method, compared with the gold standard, where differential abundance analysis is conducted with LinDA and LDM-CLR. Similarly to Supp. Fig. 5 and Supp. Fig. 6, QuanT produces a list of differentially abundant taxa that agrees best with the gold standard, followed by SVA and RUV. No correction yields the worst results when there are batch effects.

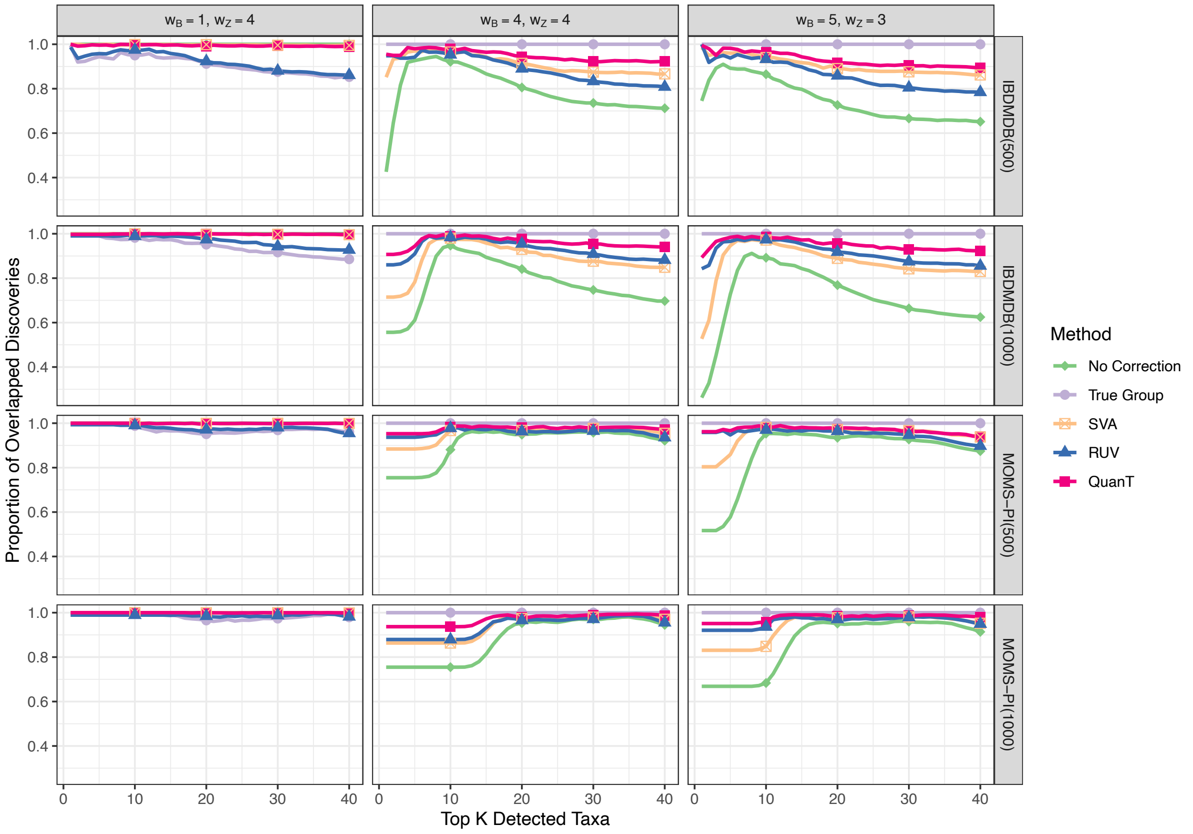


**Supp. Fig. 15 | Proportions of overlapping discoveries among the top K differentially abundant taxa identified by each method, compared with the gold standard when data are simulated from the Dirichlet-Multinomial distribution using the IBDMDB and MOMS-PI datasets as the template and analyzed with LinDA. $w_B$ and $w_z$ define the underlying batch effect and clinical effect. When there is no batch effect ($w_B = 0$), the method with no correction is the gold standard. When batch effect exists ($w_B \neq 0$), the method with true batch correction is the gold standard.**

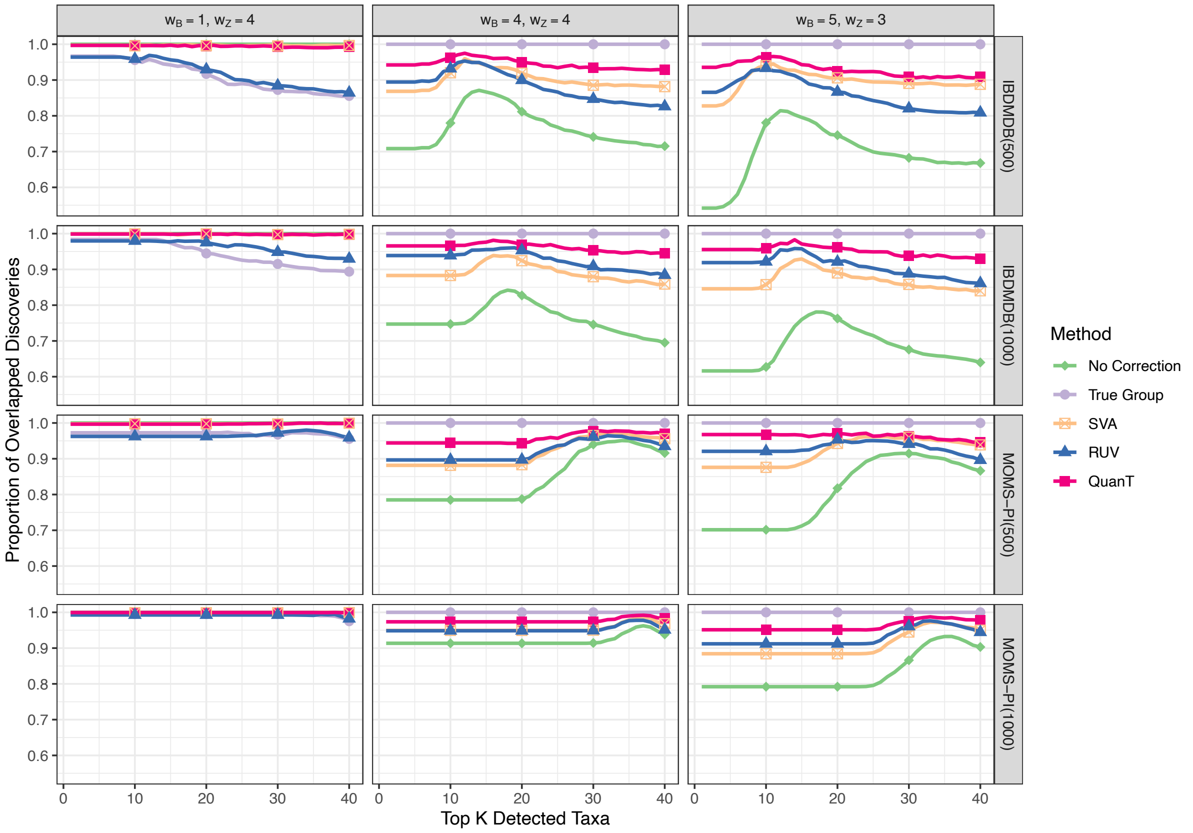


**Supp. Fig. 16 | Proportions of overlapping discoveries among the top K differentially abundant taxa identified by each method, compared with the gold standard when data are simulated from the Dirichlet-Multinomial distribution using the IBDMDB and MOMS-PI datasets as the template and analyzed with LDM-CLR. $w_B$ and $w_z$ define the underlying batch effect and clinical effect. When there is no batch effect ($w_B = 0$), the method with no correction is the gold standard. When batch effect exists ($w_B \neq 0$), the method with true batch correction is the gold standard.**

### Summary

The results based on either adjusted differential abundance analysis or batch correction analysis via ConQuR demonstrate that using QuanT's surrogate variables can lead to reliable results in downstream analysis using either approach. Using QuanT in either way has FDR well-controlled, preserves the effects of the variables of interest, removes the confounding effects of unmeasured variables, and leads to results that are the most similar to adjusting for the true hidden variables.
